# Structural Basis of Condensin Recruitment for X Chromosome Repression

**DOI:** 10.64898/2026.03.13.711519

**Authors:** Antonio Valdes Gutierrez, Gurumoorthy Amudhan, Dario Bernasconi, Sinem Erkan, Markus Hassler, Iris Suter, Brigitta Wilde, Julian Bender, Bettina Warscheid, Peter Meister, Christian H. Haering

## Abstract

In *Caenorhabditis elegans*, the condensin I^DC^ complex represses transcription from both X chromosomes in hermaphrodites to achieve dosage compensation. How condensin I^DC^ is specifically recruited to the X chromosomes in coordination with sex determination and dosage compensation (SDC) proteins and how it modulates gene expression have, however, remained unresolved. Here, we identify SDC-3 as the key adaptor that directly binds the ‘elbow’ coiled-coil domain of the condensin I^DC^-specific SMC subunit DPY-27. Using cryo-electron microscopy, we determine the structure of the SDC-3 adaptor domain bound to an auto-inhibited condensin I^DC^ holoenzyme. Disrupting this interaction compromises dosage compensation and diminishes condensin I^DC^ enrichment on the X chromosomes. Upon overcoming auto-inhibition, condensin I^DC^ exhibits robust DNA loop-extrusion activity comparable to that of canonical condensin. We propose that SDC-3-anchored condensin I^DC^ exploits loop-extrusion to reorganize X-chromosome chromatin and mediate chromosome-wide transcriptional repression.

## Introduction

Dosage compensation is an essential process that equalizes the expression of X-linked genes between the sexes when they differ in the number of sex chromosomes. In *Caenorhabditis elegans*, XX hermaphrodites achieve dosage compensation by reducing by half X-linked gene expression on both X chromosomes, thereby matching the expression levels of X0 males. Genetic screens for mutants exhibiting the hermaphrodite-specific ‘Dumpy’ phenotype (Dpy; short and fat) led to the identification and characterization of the Dosage Compensation Complex (DCC), a multi-component assembly that binds both X chromosomes to mediate transcriptional downregulation^1^.

The DCC consists of two main functional modules: (1) a group of SDC proteins—comprising SDC-1, SDC-2, SDC-3, DPY-21, and DPY-30—and (2) a specialized Structural Maintenance of Chromosomes (SMC) complex, known as condensin I^DC^. Specific recruitment of the DCC to the X chromosomes depends on recruitment elements on X (*rex*) sites distributed along the chromosome, which contain a cluster of one or more defined MEX DNA sequence motifs^2,3^. The binding partners of these sequence motifs remain uncharacterized.

In addition to SDC-2, which is thought to function as the primary determinant of X-chromosome specificity, SDC-3 and DPY-30 are required for the stable association of the entire complex, including condensin I^DC^, at *rex* sites^4^. The DCC organizes the X chromosomes in a condensin I^DC^-dependent manner into discrete ∼1 megabase-pair (Mbp) Topologically Associating Domains (TADs), a feature not observed on autosomes^5^. Whereas *rex* sites colocalize with TAD boundaries in wild-type animals, loss of condensin I^DC^ function causes clustering of multiple *rex* sites and upregulation of X-linked gene expression. Condensin I^DC^ regulates therefore both chromosome structure and gene expression^6^.

Like all eukaryotic members of the SMC family of protein complexes, condensin I^DC^ is composed of two ∼50-nm-long coiled-coil SMC subunits (MIX-1 and DPY-27), which heterodimerize via globular ‘hinge’ domains at one end of the coils and expose a pair of globular ATPase ‘head’ domains at the other end of the coils. A flexible kleisin subunit (DPY-26) bridges the heads and recruits two subunits that are almost entirely composed of α-helical HEAT (Huntingtin, Elongation factor 3, protein phosphatase 2A,TOR1)-repeat motifs (DPY-28, CAPG-1)^7^. Condensin I^DC^ differs from canonical *C. elegans* condensin I by the substitution of the SMC subunit SMC-4 with DPY-27^8^.

Condensin complexes organize the three-dimensional structure of chromosomes by extruding large DNA loops—an activity that can be inferred from mapping of mitotic chromosome architecture in cells and that can be directly visualized *in vitro* in single-molecule experiments^9–13^. Most metazoans possess two condensin complexes—condensin I and condensin II—which function in mitotic and meiotic chromosome folding and segregation. In *C. elegans*, condensin I also localizes to the interphase nucleus, where it forms long-range chromatin loops spanning 100 kilobase pairs (kbp) to 3 Mbp, effectively substituting for mammalian cohesin in shaping interphase chromatin^6^. Whether *C. elegans* condensin I—and, more intriguingly, condensin I^DC^—can extrude DNA loops has not yet been directly tested.

It has long been proposed that condensin I^DC^ is targeted to the X chromosomes through an interaction of the only condensin I^DC^-specific subunit DPY-27 with an X-linked binding partner, possibly one or more SDC proteins, and that it organizes X chromosomes by extruding DNA loops. Here, we identify the link between condensin I^DC^ and the other DCC components by mapping the interface between its DPY-27 subunit and the carboxy terminus of SDC-3 (SDC-3_C_). Disruption of the SDC-3_C_ binding interface by mutation redirects condensin I^DC^ from X chromosomes to all chromosomes and results in nematodes with a Dpy phenotype, underscoring its functional importance for dosage compensation. The cryo-electron microscopy (cryo-EM) structure of the SDC-3_C_–condensin I^DC^ holocomplex reveals an auto-inhibited state, presumably primed for DNA loading. Upon activation, condensin I^DC^ extrudes DNA loops similar to other SMC complexes. Together, these findings provide fundamental insights into how chromosome-specific targeting repurposes a conserved DNA loop extrusion machinery for gene regulation.

## Results

### Condensin I^DC^ binds and extrudes DNA

To test whether the specialized function of condensin I^DC^ in X chromosome dosage compensation is reflected by its ability to extrude DNA loops, we recombinantly expressed and purified the ∼750-kDa *C. elegans* condensin I^DC^ and condensin I holocomplexes to homogeneity (Fig. 1a). Addition of condensin I^DC^ to ∼50-kbp λ-phage DNA molecules attached to a passivated surface in a microfluidic flow chamber and fluorescently labelled with Sytox Orange resulted in the rapid formation and growth of dense spots of DNA (Fig. 1b) that could be stretched into loops by increasing buffer flow (Supplementary Fig. 1a). We never observed DNA loop extrusion in the absence of ATP or when using condensin I^DC^ complexes in which the ATPase domains had been mutated in their Walker B motifs.

**Figure 1.**
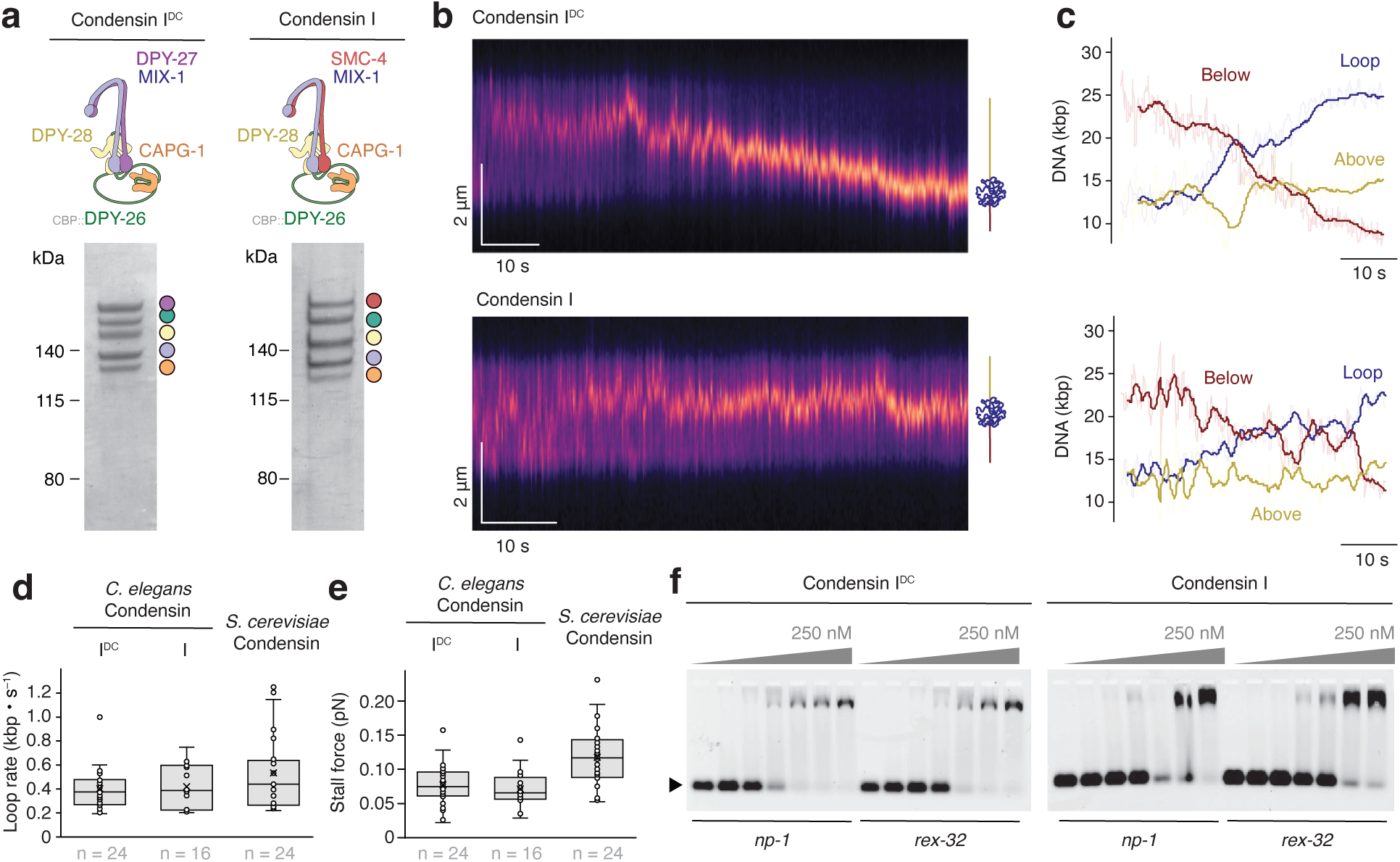
Condensin I^DC^ binds and extrudes DNA into loops. **a**, Cartoon models and Coomassie Blue-stained SDS PAGE of recombinantly expressed and purified *C. elegans* condensin I^DC^ (left) or condensin I (right). CBP: Calmodulin-Binding Peptide. **b**, Kymographs of loop extrusion on SYBR-green labeled ∼50-kbp lambda phage DNA in the presence of condensin I^DC^ (top) or condensin I (bottom) and ATP. **c**, Quantitation of fluorescence signal outside (top/bottom) or inside (loop) the DNA loops normalized to kilo-base pairs. Comparison of **d**, DNA loop extrusion rates and **e**, stalling forces of *C. elegans* condensin I^DC^ to *C. elegans* condensin I or *S. cerevisiae* condensin. The box shows the interquartile range (Q1 to Q3) with a line for the median and a cross for the mean; whiskers show minimum and maximum values within the 1.5× interquartile range. **f**, Electrophoretic mobility shift assay (EMSA) of 51-bp dsDNA substrates (10 nM) with or without a *rex* site and increasing amounts of condensin I^DC^ (left) or condensin I (right).

Analysis of fluorescence kymographs showed that condensin I^DC^—like all SMC complexes^13,14^—extruded DNA asymmetrically: the increase of the dot-like accumulated fluorescence signal (the loop) occurred simultaneously with a decrease in fluorescence on only one side of the dot signal (Fig. 1c). We obtained similar results for *C. elegans* condensin I. Average DNA loop extrusion rates of *C. elegans* condensin I^DC^ (0.40 ± 0.17 kbp·s^−1^, mean ± SD) were comparable to those of *C. elegans* condensin I (0.42 ± 0.19 kbp·s^−1^) or *S. cerevisiae* condensin (0.53 ± 0.31 kbp·s^−1^) under the same experimental conditions (Fig. 1d). Similarly, stall forces of *C. elegans* condensin I^DC^ (0.08 ± 0.03 pN, mean ± SD) were in the same range as those of *C. elegans* condensin I (0.07 ± 0.03 pN) or *S. cerevisiae* condensin (0.12 ± 0.04 pN) (Fig. 1e).

The basal ATPase rates measured in the absence of DNA for *C. elegans* condensin I^DC^ (0.8 ± 0.1 s^−1^, mean ± SD) and *C. elegans* condensin I (1.2 ± 0.4 s^−1^) were similar to the ATPase rate of *S. cerevisiae* condensin (0.5 ± 0.2 s^−1^). Addition of double-stranded DNA, however, had no effect on the ATPase activity of either *C. elegans* condensin, while it increased the ATPase rate of *S. cerevisiae* condensin more than five-fold (3.0 ± 1.0 s^−1^) (Supplementary Fig. 1b). This raises the possibility that *C. elegans* condensin I^DC^ and I may not effectively bind DNA under *in vitro* conditions.

We therefore directly assayed DNA binding properties of purified *C. elegans* condensin I^DC^ and I using double-stranded DNA substrates that either included the strong *rex* site *rex-32*, which contains both MEX and MEX II motifs and recruits the DCC *in vivo*, or a sequence from the X chromosome that is devoid of DCC binding *in vivo* (*np-1*)^15^. Both complexes bound DNA with similar apparent nanomolar affinity in electrophoretic mobility shift assays (EMSA), regardless of DNA sequence composition (Fig. 1f). In summary, our experiments demonstrate that condensin I^DC^ can bind DNA double helices independently of sequence and processively extrude them into loops, in a manner comparable to other SMC protein complexes.

### Condensin I^DC^ associates with all chromosomes in the absence of the SDC

To re-examine how condensin I^DC^ associates with chromosomes *in vivo*, we endogenously tagged the condensin I^DC^-specific subunit DPY-27 with the fluorescent protein mStayGold (Supplementary Tables 1,2). During early embryogenesis, DPY-27 fluorescence was diffusely distributed throughout the nucleus (Fig. 2a). In hermaphrodite animals, as development progressed, the DPY-27 signal became progressively restricted to one or two distinct nuclear domains, which were previously shown to be the X chromosomes using immunofluorescence microscopy and chromatin immunoprecipitation^8,16^. This transition occurred at approximately the 50-cell stage and coincided with the onset of SDC-2 protein expression, consistent with previous observations from immunofluorescence staining^8^. In contrast, DPY-27 fluorescence remained diffuse in the nuclei of male embryos at the same developmental stage (Fig. 2b).

**Figure 2.**
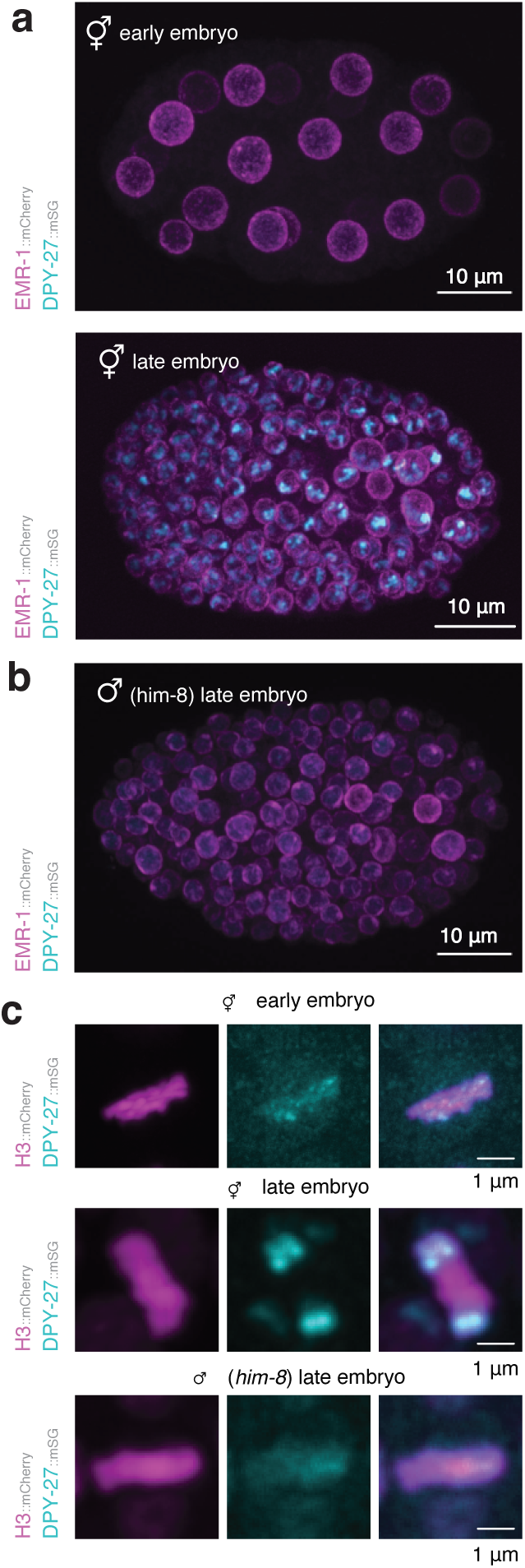
Condensin I^DC^ binds all chromosomes in the absence of the SDC. **a**, Representative confocal microscopy images of early (≤50 nuclei) or late (>50 nuclei) *C. elegans* hermaphrodite embryos expressing mCherry-labeled EMR-1 as marker for the nuclear envelope and the condensin I^DC^-specific subunit DPY-27 fused to mStayGold. **b**, Representative image of late male embryos as in panel a. **c,** Representative images of DPY-27 fused to mStayGold on metaphase chromosomes labeled with histone H3 fused to mCherry.

To determine whether the diffuse nuclear DPY-27 signal reflects chromosome-bound condensin I^DC^, we analyzed mitotic cells with their chromosomes aligned on the metaphase plate (Fig. 2c). Before the 50-cells stage, the DPY-27 signal was distributed across all chromosomes in hermaphrodites but later became restricted to one or two puncta, likely representing X sister chromatids. In male embryos, in which dosage compensation is not activated, DPY-27 continued to decorate all chromosomes at later developmental stages.

Together, the *in vivo* and *in vitro* results suggest that condensin I^DC^ associates broadly with chromosomal DNA without detectable specificity for X-linked *rex* target sites in the absence of SDC components and that its enrichment for the X chromosomes *in vivo* relies on other subunits of the DCC complex.

### SDC-3 binds the elbow region of the DPY-27 SMC coiled coil

Since our *in-vitro* data shows that *rex* sites are not directly recognized by condensin I^DC^, the latter must be targeted to the X chromosomes by one or more of the SDC proteins^1^. We therefore searched for potential DNA-binding motifs among the five SDC proteins. SDC-1 and SDC-3 contain predicted zinc finger (ZnF) motifs—domains commonly associated with sequence-specific DNA binding. Because SDC-1 is dispensable for condensin I^DC^ enrichment on the X chromosomes *in vivo*, we focused on the tandem ZnF motifs located near the carboxy terminus of SDC-3. Deletion of this region (*sdc-3(y128)*) causes severe dosage compensation defects *in vivo*^17^.

To test whether the SDC-3 ZnF motifs bind DNA, we expressed and purified the carboxy-terminal region of SDC-3 (SDC-3_C_, residues 1,764–2,150; Supplementary Fig. 2a) fused to the fluorescent protein mCherry and performed EMSA using the same conditions in which condensin I^DC^ and I showed nanomolar affinity to DNA. SDC-3_C_ failed to bind double-stranded DNA regardless of whether the DNA contained a *rex* site or not, and even at protein concentrations up to 0.5 µM (Supplementary Fig. 2b).

Because ZnF motifs also mediate protein–protein interactions^18^, we tested whether SDC-3_C_ would interact with either or both condensin I complexes by mixing mNeonGreen (mNG)-tagged SDC-3_C_ with either condensin I^DC^ or I and performing anti-mNG pulldown. SDC-3_C_ robustly and specifically co-precipitated condensin I^DC^, but not condensin I (Fig. 3a). In reciprocal co-immunoprecipitation reactions, condensin I^DC^ efficiently co-precipitated SDC-3_C_ (Supplementary Fig. 2c). We furthermore validated complex formation between mNG-tagged SDC-3_C_ and condensin I^DC^ by analytical size exclusion chromatography (SEC) (Fig. 3b). In contrast, we observed no interaction between condensin I^DC^ and the other two SDC proteins DPY-21 and DPY-30 (Supplementary Fig. 2c). Neither SDC-3_C_, DPY-21, nor DPY-30 affected the ATPase activity of condensin I^DC^ when added in equimolar amounts (Supplementary Fig. 2d). Similarly, addition of SDC-3_C_ had no significant effect on either DNA loop extrusion rates (0.4 ± 0.2 kbp·s^−1^ –SDC-3_C_, 0.5 ± 0.2 kbp·s^−1^ +SDC-3_C_, mean ± SD) or stall forces (0.07 ± 0.03 pN –SDC-3_C_, 0.08 ± 0.03 pN +SDC-3_C_; mean ± SD) of condensin I^DC^ (Supplementary Fig. 2e). To further characterize the interface between SDC-3_C_ and condensin I^DC^, we performed crosslinking mass spectrometry (XL-MS) using the bifunctional crosslinker BS3 (Bis(sulfosuccinimidyl)suberate) that predominantly reacts with primary amines but also displays reactivity towards hydroxyl groups^19^. As expected, XL-MS identified numerous intra-and inter-subunit cross-links within the condensin I^DC^ complex, as well as a single crosslink between SDC-3_C_ and condensin I^DC^ (Fig. 3c, Supplementary Table 3). This crosslink connects lysine residues of SDC-3_C_ and a region of the DPY-27 coiled coil that corresponds to the conserved ‘elbow’ region characteristic of coiled-coil dimers in multiple SMC complexes (Supplementary Fig. 2f)^20,21^.

**Figure 3.**
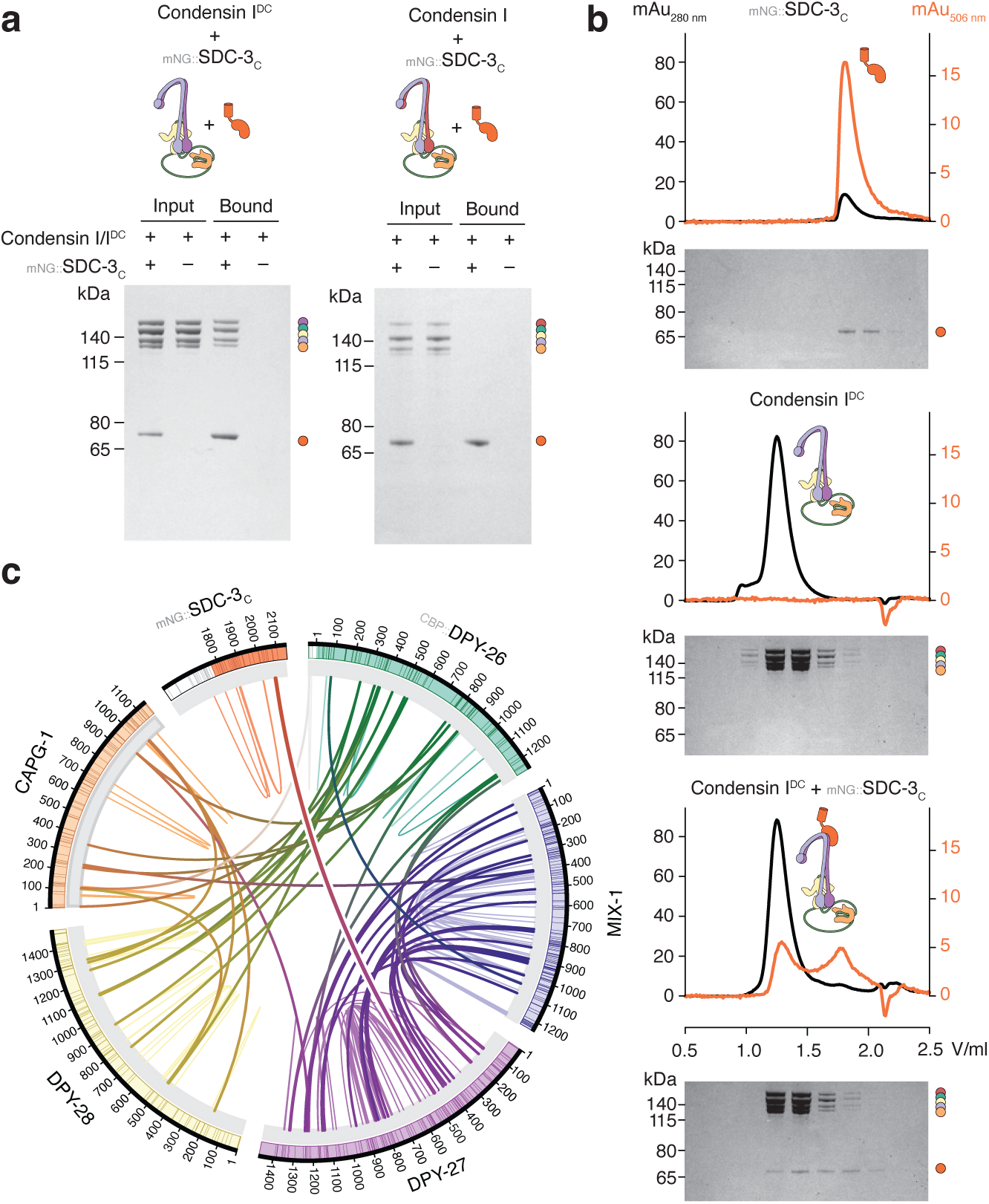
The carboxy terminus of SDC-3 binds condensin I^DC^. **a**, Coomassie Blue-stained SDS PAGE of condensin I^DC^ (left) or condensin I (right) input and bound fractions of immunoprecipitation reactions against SDC-3_C_ fused to mNeonGreen (mNG). **b**, Analytical SEC profiles for total protein absorption (280 nm, black) or mNG (506 nm, orange) and Coomassie Blue-stained SDS PAGE of mNG::SDC-3_C_ (top), condensin I^DC^ (middle), or an equimolar mix of both (bottom). **c**, Circos plot of intra- (thin lines) or inter-molecular (thick lines) crosslinks identified in condensin I^DC^ holocomplexes bound to SDC-3_C_.

To test whether the DPY-27–MIX-1 elbow of condensin I^DC^ was sufficient for SDC-3_C_ binding, we expressed and purified a truncated DPY-27–MIX-1 dimer that contained only elbow coiled coil and hinge dimerization domains. Analytical SEC showed that the DPY-27–MIX-1_elbow_ dimer formed a stable complex with SDC-3_C_ (Fig. 4a) with similar efficiency as it did with the condensin I^DC^ holocomplex (Fig. 3b). XL-MS identified two additional cross-links between SDC-3_C_ and the DPY-27–MIX-1_elbow_ dimer, expanding upon the single link detected with the condensin I^DC^ holocomplex (Supplementary Fig. 2g and Supplementary Table 4).

**Figure 4.**
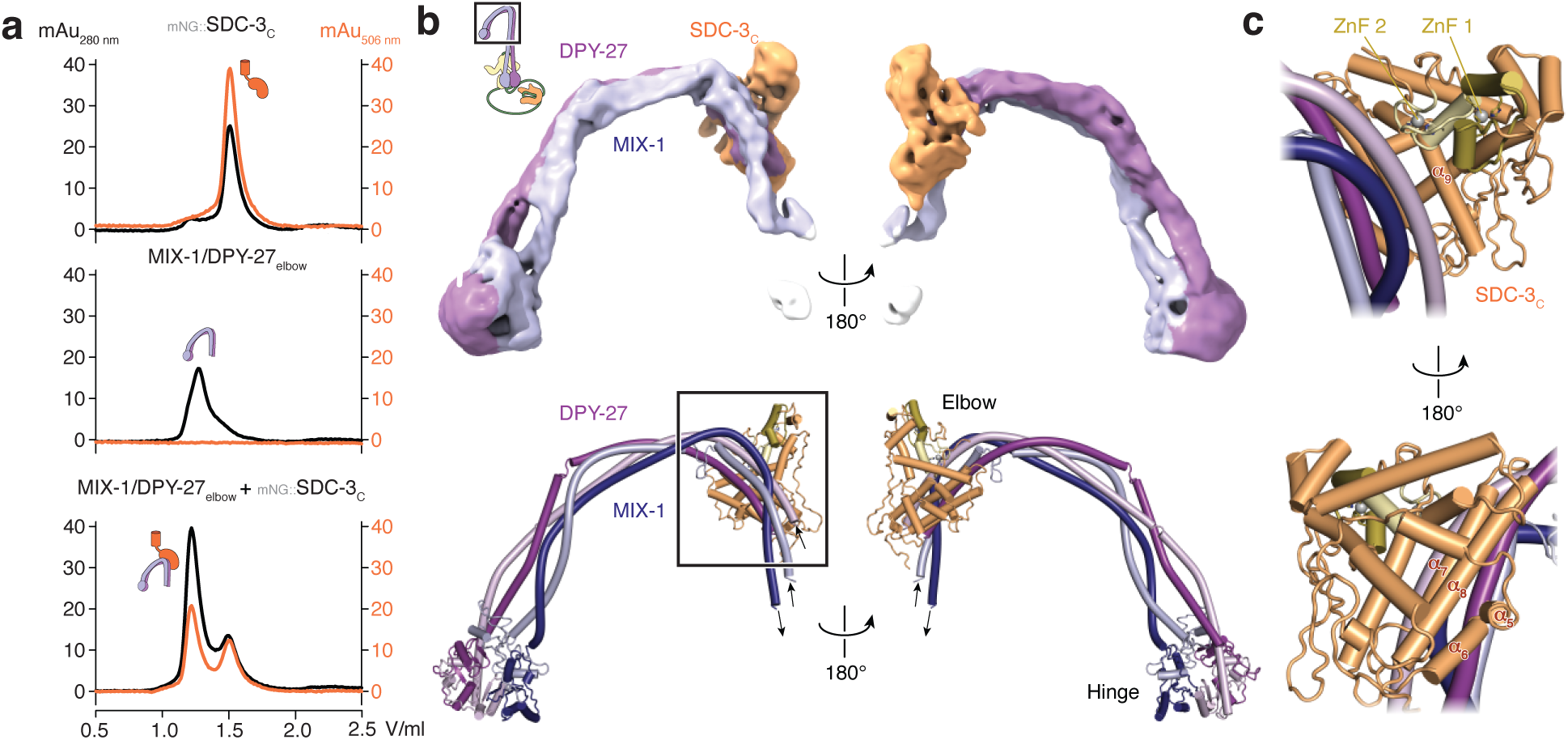
SDC-3_C_ binds the coiled-coil elbow region of DPY-27. **a**, Analytical SEC profiles for mNG::SDC-3_C_ (top), the MIX-1–DPY-1_elbow_ dimer (middle), or an equimolar mix of both (bottom). **b**, Cryo-EM density map (top) and model (bottom) of SDC-3_C_ (orange) bound to the MIX-1–DPY-1_elbow_ dimer (blue–purple). **c**, Close-up views of the SDC3_C_–DPY-27 interface (box in (B)) highlight the SDC-3_C_ helices that form a bundle with the DPY-27 coiled coil and the SDC-3_C_ zinc finger motifs (ZnF 1/2).

### Cryo-EM structure of the SDC-3_C_ interface with DPY-27

We elucidated the molecular basis of the interaction between SDC-3_C_ and the condensin I^DC^ holocomplex by determining their co-structure using cryo-EM. To stabilize condensin I^DC^ in a nucleotide-engaged state suitable for single-particle analysis, we used an ATP hydrolysis-deficient ‘Walker B’ mutant version (DPY-27_E1274Q_– MIX-1_E1129Q_) in the presence of adenosine triphosphate (ATP). Despite this stabilization, electron micrographs revealed substantial conformational variability in the SMC coiled-coil domains, similar to what had previously been observed for *S. cerevisiae* condensin^13,22^. We therefore processed particle images of the coiled coils containing the elbow and the hinge regions separately from the catalytic SMC core bound to the kleisin and HEAT-repeat subunits (Supplementary Fig. 3).

Processing of particles yielded a 7.0 Å-resolution reconstruction of the DPY-27–MIX-1 elbow region (Supplementary Fig. 4 and Table 1). The map displayed additional density at the bend of the coiled coil that could not be assigned to either SMC subunits. Guided by AlphaFold3^23^, we fitted a model of SDC-3_C_ into this unassigned density (Fig. 4b). The vast majority (114 of 119) of crosslinks identified by XL-MS for the DPY-27–MIX-1_elbow_ dimer bound to SDC-3_C_ connect residues within distances compatible with the BS3 crosslinker, thereby confirming the structural model (Supplementary Fig. 5a).

**Table 1.** Cryo-EM data collection, processing, and model statistics.

|  | Condensin I<sup>DC</sup><br>core complex<br>PDB ID **9TZA**<br>**EMD-56463** | Condensin I<sup>DC</sup><sub>elbow</sub>–<br>SDC3<sub>C</sub> complex<br>PDB ID **9TZB**<br>**EMD-56464** | Condensin I<sup>DC</sup><br>core dimer<br>**EMD-56465** |
| --- | --- | --- | --- |
| **Data collection and Processing** |  |  |  |
| Microscope | Titan Krios G3 | Titan Krios G3 | Titan Krios G3 |
| Voltage (keV) | 300 | 300 | 300 |
| Camera | Falcon IVi | Falcon IVi | Falcon IVi |
| Magnification | ×130,000 | ×130,000 | ×130,000 |
| Pixel size at detector (Å/pixel) | 0.946 | 0.946 | 0.946 |
| Total electron exposure (e<sup>-</sup>/Å<sup>2</sup>) | 70 | 70 | 70 |
| Exposure rate (e<sup>-</sup>/pixel/sec) | 10.13 | 10.13 | 10.13 |
| Number of frames collected during exposure | 1800 | 1800 | 1800 |
| Defocus range (μm) | –1.2 to –3.0 | –1.2 to –3.0 | –1.2 to –3.0 |
| Automation software | EPU 3.8.1. | EPU 3.8.1. | EPU 3.8.1. |
| Energy filter slit width (eV) | 5.0 | 5.0 | 5.0 |
| Micrographs collected/used (no.) | 14,940/14,900 | 14,940/14,900 | 7,221/7,208 |
| Total extracted particles (no.) | 3,066,606 | 397,855 | 631,654 |
| Final particle images (no.) | 202,339 | 80,318 | 4,432 |
| Symmetry imposed | C1 | C1 | C1 |
| Map Resolution (global, Å) |  |  |  |
| FSC 0.143 (unmasked/masked) | 4.10/3.34 | 8.40/6.96 | 18.50/8.73 |
| Resolution range (local, Å) | 3.1 – 8.3 | 6.9 – 12.7 | N/A |
| Map sharpening *B* factor (Å<sup>2</sup>) | –71 | –588 | –586 |
| **Refinement** |  |  |  |
| Refinement package | PHENIX 1.21-5419 | PHENIX 1.21-5419 | N/A |
| Real space or geometry minimization | Real space | Geometry minimization |  |
| Model composition |  |  |  |
| Non-hydrogen atoms | 34,686 | 11,505 |  |
| Protein | 4,319 | 1,428 |  |
| Ligands | 2 | 2 |  |
| Model-Map scores |  |  |  |
| Overall correlation coefficients |  |  |  |
| CC (mask) | 0.79 | 0.36 |  |
| CC (volume) | 0.78 | 0.35 |  |
| Map versus Model FSC 0.143 (Å) | 3.32 | N/A |  |
| Average *B* factors (Å<sup>2</sup>) |  |  |  |
| Protein | 174 | N/A |  |
| Ligand | 121 | N/A |  |
| R.m.s. deviations from ideal values |  |  |  |
| Bond lengths (Å) | 0.004 | 0.004 |  |
| Bond angles (°) | 0.788 | 0.845 |  |
| Validation |  |  |  |
| MolProbity score | 1.55 | 2.03 |  |
| CaBLAM outliers (%) | 1.70 | 2.54 |  |
| Clashscore | 6.93 | 6.81 |  |
| Poor rotamers (%) | 0.31 | 2.12 |  |
| Ramachandran plot |  |  |  |
| Favored (%) | 97.05 | 93.80 |  |
| Allowed (%) | 2.95 | 5.41 |  |
| Outliers (%) | 0.00 | 0.70 |  |

SDC-3_C_ adopts a predominantly α-helical globular fold with no close structure homologs detected in a DALI search^24^. Both Cys_2_His_2_ ZnF motifs align along one face of the protein, each coordinating a zinc ion (Supplementary Fig. 4c). Extensive intra-molecular contacts between the SDC-3_C_ ZnF motifs occlude potential DNA binding surfaces, which most likely explains the lack of DNA binding in EMSA (Supplementary Fig. 2b). The primary interface with condensin I^DC^ is mediated by helices α_6_ and α_7_, as well as the second of the two ZnF motifs of SDC-3_C_ (Fig. 4c,5a), which contact exclusively the amino- and carboxy-terminal helices of the DPY-27 coiled coil. In addition to their role in stabilizing the global fold of the SDC-3_C_ domain, the contribution of one of the ZnF motifs to the binding interface explains why their deletion disrupts condensin I^DC^ X chromosome localization^25^. The specific interaction with DPY-27 explains the selective recruitment to X chromosomes of condensin I^DC^—but not of condensin I, which contains SMC-4 in place of DPY-27—by the SDC proteins.

**Figure 5.**
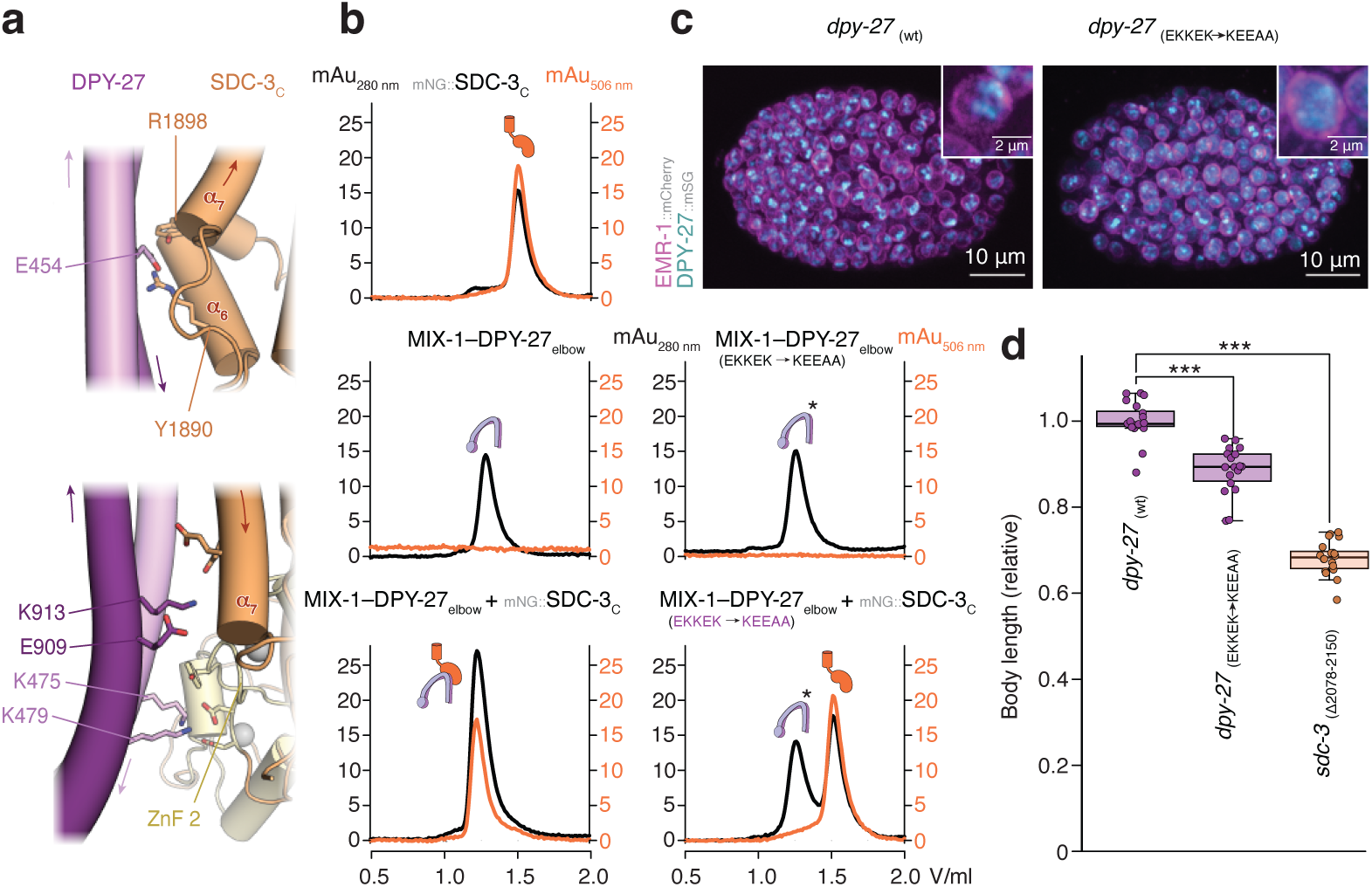
DPY-27 mutations that disrupt SDC-3_C_ binding cause a Dpy phenotype in *C. elegans* animals. **a**, Close-up views of the DPY-27_elbow_–SDC-3_C_ interface highlighting residues selected for mutagenesis. **b**, Analytical SEC profiles for mNG::SDC-3_C_ (top), unmodified (left) or E_454_K, K_475_E, K_479_E, E_909_A, K_913_A mutant version (right) of the MIX-1–DPY-27_elbow_ dimer, or an equimolar mixture of both proteins. **c**, Representative confocal microscopy images of wild-type DPY-27 (left) or homozygous E_454_K, K_475_E, K_479_E, E_909_A, K_913_A DPY-27 mutant (right) fused to mStayGold in late hermaphrodite embryos expressing mCherry-labeled EMR-1 as marker for the nuclear envelope. **d**, Relative body lengths of *dpy-27::mSG* wild-type (strain PMW1297), homozygous *dpy-27::mSG* E_454_K, K_475_E, K_479_E, E_909_A, K_913_A mutant (strain PWM1373), or *sdc-3(Δ2078–2150)* mutant (strain PMW1431) adult worms. The box shows the interquartile range (Q1 to Q3) with a line for the median; whiskers show minimum and maximum values within the 1.5× interquartile range (*** p < 0.001, t-test or Wilcoxon rank-sum test, depending on normality and variance assumptions, with p-values adjusted for multiple comparisons using the FDR method).

### Disruption of the SDC-3_C_–DPY-27 interface results in a Dpy phenotype

To probe the functional relevance of the SDC-3_C_–DPY-27 interaction identified in our structural model, we simultaneously mutated charged residues in the DPY-27 coiled coil at the interface with SDC-3_C_ to residues of opposite charge: three conserved residues in the amino-terminal helix (E_454_, K_475_, and K_479_) and two charged residues in its carboxy-terminal helix (E_909_ and K_913_; Fig. 5a, Supplementary Fig. 5b). The mutations in DPY-27 abolished SDC-3_C_ co-elution with the MIX-1–DPY-27_elbow_ dimer during analytical SEC (Fig. 5b) and prevented co-immunoprecipitation of SDC-3_C_ with both condensin I^DC^ holocomplexes (Supplementary Fig. 5c) or MIX-1–DPY-27_elbow_ dimers (Supplementary Fig. 5d).

Introduction of the interface mutations into the endogenous alleles of *C. elegans* resulted in a diffuse localization of DPY-27 in the nuclei of hermaphrodite embryos (Fig. 5c, Supplementary Fig. 5e), which suggests that SDC-3_C_-binding-deficient mutant condensin I^DC^ fails to target uniquely the X chromosomes. As these embryos developed into adults, they displayed a Dpy phenotype, with their body length significantly reduced compared to DPY-27 wild-type animals; although the effect was less severe than the Dpy phenotype observed in animals lacking the entire SDC-3_C_ region (Fig. 5d).

Identifying residues in SDC-3_C_ that mediate key interactions with the DPY-27 coiled coil was challenging due to the low resolution of the interface cryo-EM density map and the lack of alternative structural information. Simultaneous mutation of two conserved residues (Y_1890_ and R_1898_; Supplementary Fig. 5f) in SDC-3_C_ helix α_6_ abolished co-immunoprecipitation of condensin I^DC^ holocomplexes (Supplementary Fig. 5g) and co-elution with the MIX-1–DPY-27_elbow_ dimer during analytical SEC (Supplementary Fig. 5h). However, DPY-27 still formed clusters in hermaphrodite animals expressing only the mutant version of SDC-3_C_ (Supplementary Fig. 5i), and these animals developed to near wild-type length—unlike those carrying both the two point mutations and deletions of the two ZnF motifs, which exhibited a clear Dpy phenotype (Supplementary Fig. 5j). These results suggest that, while the SDC-3_C_–DPY-27 interface plays a functionally important role in recruiting condensin I^DC^, additional interactions may also contribute to its recruitment to or stabilization on X chromosomes.

### Cryo-EM structure of an auto-inhibited condensin I^DC^ catalytic core

Processing of particles of the condensin I^DC^ core complex (Supplementary Fig. 4) resulted in a 3.3 Å-resolution electron density map of the nucleotide-engaged MIX-1–DPY-27 ATPase head dimer with adjacent coiled coils, the majority of the DPY-26 kleisin subunit, and both HEAT-repeat subunits (Fig. 6a). The quality of the map allowed *de novo* building of a structural model of the entire core complex (Supplementary Fig. 6a, Table 1), which revealed an overall conformation that had not previously been described for SMC protein complexes: The HEAT-A subunit DPY-28 packs onto one side of the MIX-1–DPY-27 dimer and recruits the HEAT-B subunit CAPG-1 to the complex (Fig. 6b). The DPY-26 kleisin subunit, which is substantially longer than any previously structurally characterized kleisin, serves as a central organizing scaffold, forming contacts with all four remaining subunits and effectively holding the complex together.

**Figure 6.**
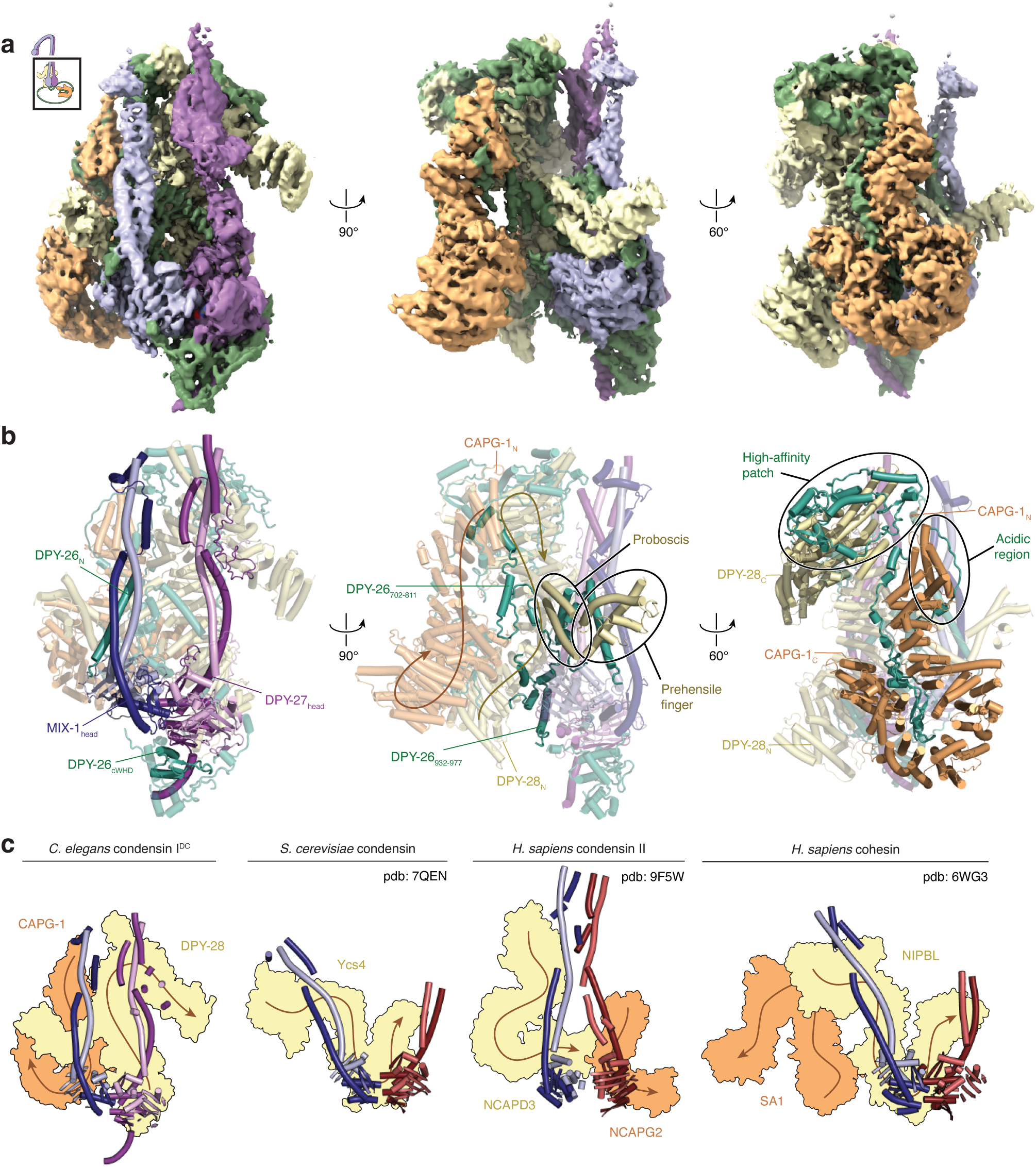
Cryo-EM structure of the *C. elegans* condensin I^DC^ core complex. **a**, Electron density map and **b**, model of the *C. elegans* condensin I^DC^ core complex. The orientation of the DPY-28 HEAT-A subunit (yellow) and CAPG-1 HEAT-B subunit (orange) packed via interactions with the DPY-26 kleisin subunit (green) onto the MIX-1 (blue) and DPY-27 (purple) SMC subunits is indicated by arrows. A ‘prehensile finger’ extension at the tip of the DPY-28 proboscis, a large DPY-26 kleisin extension located at the ‘high affinity patch’ binding site of DPY-28, and an acidic DPY-26 kleisin region that occupies the DNA binding site in CAPG-1 are highlighted. **c**, Orientation of the HEAT-A (yellow) and HEAT-B (orange) subunits of the *C. elegans* condensin I^DC^ structure (left) compared to structures of *S. cerevisiae* condensin, human condensin II, or human cohesin.

In previously determined condensin or cohesin structures, the HEAT-A subunit interacts with the engaged ATPase heads in an orientation that follows their U-shaped arrangement, with its amino-terminal ‘leg’ aligned with the ν-SMC coiled coil and its carboxy-terminal ‘leg’ aligned with the κ-SMC coiled coil^13,20,26–29^. In our condensin I^DC^ core structure, the orientation of the HEAT-A subunit DPY-28 is flipped (Fig. 6c), effectively blocking the trajectory of a DNA double helix through the coiled-coil and on top of the nucleotide-engaged ATPase head ‘motor’ site that was observed in other condensin and cohesin ‘gripping state’ structures.

A second striking feature of DPY-28 is an additional helical domain at the tip of the so-called ‘proboscis’—an α-helical protrusion from the HEAT-repeat backbone previously reported in the structure of the homologous Ycs4 protein in *S. cerevisiae* absent in the homologous NCAPD3 HEAT-A subunit of a human condensin II structure^26,30^. Structure predictions deposited in the AlphaFold database suggests that this additional domain is a feature common to animal condensin I HEAT-A subunits (Supplementary Fig. 6b)^31^. In analogy to the tip of an elephant’s proboscis and its potential molecular function (see Discussion), we termed the additional domain ‘prehensile finger’ (Fig. 6b, Supplementary Fig. 6b). The prehensile finger wraps around the triple helix formed by the MIX-1 coiled-coil neck region and the amino-terminal DPY-26 kleisin ‘contact’ helix, presumably preventing the two short helices that are predicted to precede the contact helix from forming the helix-turn-helix motif seen in other SMC complexes. The proboscis coils are presumably stabilized by a DPY-26-specific insertion of three helices (Supplementary Fig. 7a; Ins1 α_A–C_). Furthermore, long sequence insertions in the DPY-26 kleisin at its ‘high-affinity’ binding patch for the DPY-28 HEAT-A subunit create additional contacts that presumably stabilize its U-shaped conformation (Fig. 6b, Supplementary Fig. 7a; Ins2 and Ins3)^30^.

The CAPG-1 HEAT-B subunit packs onto the DPY-28 HEAT-A subunit of condensin I^DC^, without directly contacting the SMC subunits (Fig. 6b). In yeast condensin structures, the region of the kleisin subunit that associates with the HEAT-B subunit forms a ‘safety belt’ loop structure that is closed off by the contact of ‘latch’ and ‘buckle’ motifs and topologically entraps DNA^13,32^. In the *C. elegans* condensin I^DC^ structure, the helix that most likely corresponds to the latch is disengaged and, together with a safety-belt helix that contains a highly conserved basic patch, mediates the contact to the DPY-28 HEAT-A subunit. The position of the latch helix in yeast is instead occupied by an insertion composed of two α-helices and two β-strands (Supplementary Fig. 7a; Ins4). Furthermore, the concave side of the amino-terminal ‘leg’ of the HEAT-B subunit that is occupied by DNA in the homologous yeast condensin structures is occupied by a stretch of negatively charged kleisin residues that precedes the safety belt buckle, effectively preventing DNA binding at the ‘anchor’ site of the complex. Although individual residues in this region of the ɣ-kleisin subunits of other condensin complexes are not highly conserved, several of these regions display a high degree of negative charge (Supplementary Fig. 7b). It is therefore conceivable that binding of the acidic region to the DNA anchor site at the HEAT-B subunit is a general regulatory feature (see Discussion).

Finally, DPY-26 contains three additional insertions in its carboxy-terminal quarter (Supplementary Fig. 7a): (1) A pair of anti-parallel helices that bind the DPY-27 ATPase head domain at its coiled-coil stem and mediate contacts with DPY-28 (Ins5), (2) A long insertion that is partially disordered in our structure and contacts the MIX-1 ATPase head (Ins6), and (3) an extension at the very carboxy terminus that contacts the DPY-27 ATPase head (Ins7). The additional contacts mediated by these insertions presumably stabilize the compact overall conformation of the condensin I^DC^ core.

### A minor fraction of condensin I^DC^ complexes forms dimers

Unexpectedly, single particle analysis of the cryo-EM data revealed a small population (∼3 %) of dimers (Fig. 7a, Supplementary Fig. 3b). Despite the low particle number, we were able to reconstruct an 8.7 Å-resolution map for the condensin I^DC^ dimer (Fig. 7b). The map clearly shows that two condensin I^DC^ core complexes dimerize via their CAPG-1 HEAT-B subunits facing in opposite orientations, similar to the dimers observed for the *Escherichia coli* SMC-like MukBEF complex^33^. However, the limited resolution did not allow creating a high-confidence model or selecting specific residues at the dimer interface for designing specific mutations that selectively prevent dimerization to assess their functional relevance. The dimers are clearly not an artifact of freezing complexes for cryo-EM, since we also observed a small fraction of dimers in solution by analytical SEC coupled to multi-angle light scattering (SEC-MALS; Fig. 7c) or mass photometry (Fig. 7d).

**Figure 7.**
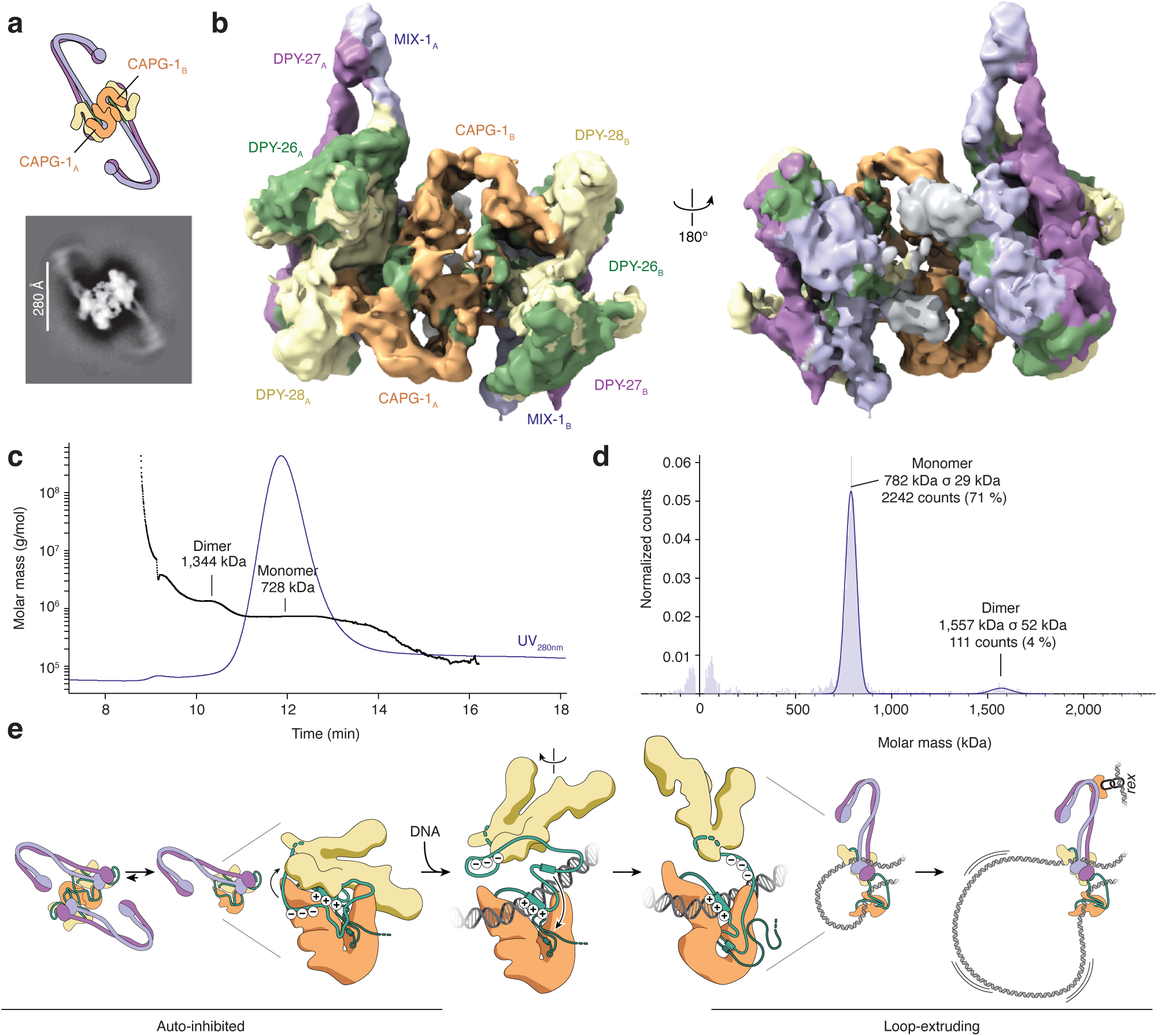
A fraction of condensin I^DC^ forms dimers. **a**, Schematic cartoon and 2D cryo-EM class average image of condensin I^DC^ holocomplex dimers. **b**, Cryo-EM density map of condensin I^DC^ core dimers. **c**, Analytical SEC coupled to multi-angle light scattering (SEC-MALS) profile of condensin I^DC^. **d**, Mass photometry profile of condensin I^DC^ (10 nM). **e** Model for the activation of auto-inhibited condensin I^DC^ for DNA loop extrusion. For details see Discussion.

## Discussion

### SDC-3 recruits Condensin I^DC^ to the X chromosomes

Whereas different SMC protein complexes are recruited to defined genomic loci through sequence-specific binding partners, such as CTCF in the case of cohesin or ParB in the case of the *Bacillus subtilis* SMC complex, condensin I^DC^ is unique in its ability to target a specific chromosome^34,35^. Although previous genetic studies established that the SDC subunits SDC-2 and SDC-3 are essential for condensin I^DC^ recruitment to the X chromosomes, the underlying targeting mechanism had remained elusive.

In this study, we identified a zinc finger-containing domain located at the carboxy terminus of SDC-3 that directly binds DPY-27, the sole condensin I^DC^-specific subunit (Fig. 3). Our cryo-EM structure reveals that two α-helices of the SDC-3_C_ form a helical bundle with the bent elbow region of the coiled coil of the κ-SMC subunit DPY-27 (Fig. 4). In this conformation, both ZnF motifs are sterically occluded from engaging a DNA double helix, consistent with their failure to bind DNA with sub-micromolar affinity (Supplementary Fig. 2b). The only other protein reported to engage an SMC coiled coil in a comparable manner is the SUMO ligase Mms21/Nse2, which forms a helical bundle with the (non-bent) coiled coil of the κ-SMC subunit Smc5^36,37^.

Mutations that disrupt the SDC-3–DPY-27 interface *in vitro* perturb, but do not completely abolish, the sharply defined localization pattern of condensin I^DC^ in late hermaphrodite embryos (Fig. 5, Supplementary Fig. 5). Consistent with dosage compensation defects, they similarly reduce hermaphrodite adult body length, although to a lesser extent than complete deletion of the SDC-3_C_ ZnF-domain. Together, these findings raise the possibility that additional condensin I^DC^-binding sites exist within full-length SDC-3 or other SDC components. Given that DPY-21 and DPY-30 do not exhibit detectable affinity for condensin I^DC^ (Supplementary Fig. 2c) and that SDC-1 is dispensable for X-chromosome targeting of condensin I^DC^ *in vivo*^38^, the most plausible candidate for additional interactions is SDC-2.

The mechanism by which SDC components recognize MEX motifs on the X chromosomes remains unclear. Among the known subunits, only SDC-1 and SDC-3 are predicted to harbor sequence-specific DNA-binding motifs. However, the two ZnF motifs in SDC-3 do not exhibit DNA-binding activity *in vitro* (Supplementary Fig. 2b), and SDC-1 is dispensable for condensin I^DC^ targeting to the X chromosomes *in vivo*^38^. These findings raise the possibility that sequence-specific DNA-binding elements may reside within SDC-2 or SDC-3 but have eluded detection by standard sequence analyses. Alternatively, additional, yet unidentified, factors may facilitate the recruitment of SDC components to their chromosomal targets.

### Condensin I^DC^ DNA loop extrusion explains DCC spreading along the X chromosomes

Our findings integrate with and extend previous models in which condensin I^DC^ itself does not directly recognize specific elements (*rex* sites) on the X chromosome^3^. Instead, SDC components first bind to these sites and then retain condensin I^DC^ at these sites *via* direct protein-protein interactions to form an integral dosage compensation complex (DCC). In this scenario, condensin I^DC^ loads stochastically along the X chromosome and autosomes, consistent with its intrinsic sequence-independent ability to bind DNA with nanomolar affinity (Fig. 1f) and its dispersed chromosomal localization in the absence of SDC components (Fig. 2). It then translocates via loop extrusion until it either dissociates or encounters SDC components at *rex* sites. Once anchored, the motor activity of condensin I^DC^, which presumably remains active when bound to SDC-3_C_ (Supplementary Fig. 2e), drives DCC spreading to occupy and repress other regions of the X chromosome.

While the first direct evidence that canonical *C. elegans* condensin I is a self-sufficient DNA loop extruder (Fig. 1) explains how it functions as an organizer of interphase autosomes in nematodes, condensin I^DC^-dependent DNA loop extrusion explains how it organizes the X chromosomes into TAD-like structures that are absent on autosomes^5,6^. Since insertion of a single *rex* site into an autosome recruits condensin I^DC^ but is insufficient to generate TADs^39^, the establishment of self-interacting chromosome domains requires boundary elements, consistent with the hypothesis that loop-extruding condensin I^DC^ complexes remain anchored at *rex* sites. Support for *rex*-mediated anchoring of condensin I^DC^ is further provided by DPY-27 Hi-ChIP data, in which contacts between *rex* sites and the TAD interior are enriched compared to Hi-C data^40^.

Because condensin I^DC^ is required not only to establish but also to maintain repression of X-linked genes^6^, continuous loop extrusion may be necessary to counteract transcriptional activity—possibly by restricting RNA polymerase II occupancy at X-linked promoters^41^. This suggests a dynamic interplay akin to the myth of Sisyphus, eternally pushing a boulder uphill only for it to roll back down: a constant tug-of-war between condensin I^DC^–mediated DNA loop extrusion and RNA polymerase–driven transcription. Rather than functioning as a binary on/off switch, this balance may produce a dynamic equilibrium, offering a mechanistic explanation for why dosage compensation reduces, but does not fully silence, X-linked gene expression.

### A primed state of the SMC core complex for DNA loading?

Our study presents the first high-resolution cryo-EM structure of a condensin I complex, revealing distinct architectural features not observed in previous structures of SMC complexes. Among the most striking is an extended projection from the ‘proboscis’ of the DPY-28 HEAT-A subunit, which we term ‘prehensile finger’ (Fig. 6). This feature is most likely present in other animal condensin I complexes (Supplementary Fig. 6b). Reminiscent of the tip of an elephant’s trunk gripping objects, the prehensile finger appears to grasp the MIX-1 coiled coil with its bound DPY-26 contact helix. This interaction may restrict ATPase head separation, analogous to the constraint imposed by binding of the Ycs4 HEAT-A amino terminus to the Smc2 neck region in the ‘bridged’ state of yeast condensin^20^.

Even more striking is the inverted orientation of the two HEAT-repeat subunits bound to the SMC ATPase core—a configuration that contrasts with those observed in previously characterized condensin and cohesin complexes (Fig. 6c)^13,20,26–29^. This novel conformation cannot be attributed to SDC-3 binding at the DPY-27 elbow region, since electron micrograph class averages from samples prepared in the absence of SDC-3_C_ exhibited the same overall architecture (Supplementary Fig. 3c). Rather, the distinctive arrangement is most likely maintained by the unusually long DPY-26 kleisin subunit, which acts as a molecular scaffold linking the subunits within the complex (Supplementary Fig. 7a).

The densely packed conformation is incompatible with DNA entrapment at the ‘motor’ site—the region between the SMC coiled coils and above the nucleotide-engaged ATPase head domains—as observed in the ATP-bound ‘gripping’ states of other SMC complexes^13,20,28,29^. Likewise, albeit the groove at the ‘anchor’ site of the CAPG-1 HEAT-B subunit is sufficiently large to accommodate a DNA double helix, positioning DNA along the same trajectory seen in yeast condensin structures^13,32^ would lead to steric clashes with a conserved acidic region of the DPY-26 kleisin subunit that occupies part of the DNA-binding interface (Supplementary Fig. 7b). It is therefore plausible that the condensin I^DC^ core structure represents an auto-inhibited conformation poised for DNA loading, which may be stabilized by the unusually long kleisin subunit of *C. elegans* but is otherwise too transient to be captured in other SMC complexes. In this scenario (Fig. 7e), DNA engagement at the anchor site would first require displacement the kleisin acidic region and then—assisted by a conserved positive patch α helix—closure of the adjacent kleisin ‘safety belt’, which is open and engages the DPY-28 HEAT-A subunit in our structure. This conformational transition likely disrupts the interactions with DPY-28, releasing the compact arrangement and activating the complex for DNA loop extrusion.

Although the ability of eukaryotic SMC complexes to function as dimers has long been discussed^12,42–45^, structural evidence for dimerization has thus far been limited to prokaryotic MukBEF and Wadjet complexes^33,46,47^. We observe that a small fraction of condensin I^DC^ complexes form dimers on cryo-EM grids, as well as in bulk and single-molecule assays (Fig. 7a–d). In these dimers, the two SMC coiled coils face in opposite directions, adopting an arrangement similar to that observed in the DNA-free structures of MukBEF^33^. While the electron density map obtained from the dimer fraction is of insufficient resolution to identify residues suitable for mutagenesis to specifically disrupt dimerization, it reveals that dimerization is mediated by the CAPG-1 HEAT-repeat subunit. Future studies will be required to assess the functional relevance of SMC complex dimerization.

## Methods

### Protein expression and purification

#### C. elegans condensin I and I^DC^ holocomplexes

Codon-optimized cDNAs encoding the five subunits of wild-type condensin I complex or wild-type or MIX-1_E1129Q_–DPY-27_E1274Q_ mutant condensin I^DC^ complexes under control of galactose-inducible promoters were integrated into the *TPR1* locus (pGAL10-DPY-28 pGAL1-CAPG-1) and the *URA3* locus (pGAL10-CBP::DPY-26 or pGAL10-HA_6_::CBP::DPY-26 pGAL1-MIX-1 pGAL7-DPY-27 or pGAL7-SMC-4) of the *S. cerevisiae* genome. Yeast cells were grown at 30 °C in YEP medium containing 2 % (w/v) D-glucose. For expression, cells were collected by centrifugation and transferred to YEP medium containing 2 % (w/v) raffinose for 6 h. Expression was induced after shifting cells to 24 °C by addition of D-galactose to 2 % (w/v) for 16 h.

Cells were collected by centrifugation, resuspended in lysis buffer (50 mM TRIS-HCl pH 7.5, 200 mM NaCl, 5 % (v/v) glycerol, 2 mM CaCl_2_, 1 mM DTT) supplemented with cOmplete EDTA-free protease inhibitor mix (cOm–EDTA, Roche), and lysed in a FreezerMill (Spex) under liquid nitrogen. The lysate was thawed on ice and cleared by 30 min centrifugation at 48,000 ×g at 4 °C before incubation with 2 mL Calmodulin Sepharose 4B beads (Cytiva) for 3 h at 4 °C. Beads were packed into a gravity flow column and washed with 30–40 column volumes (cv) lysis buffer before elution in 7–10 cv elution buffer (50 mM TRIS-HCl pH 7.5, 200 mM NaCl, 5 % (v/v) glycerol, 2 mM EGTA, 1 mM DTT).

For condensin I, the eluate from the calmodulin beads was diluted with low-salt buffer (25 mM TRIS-HCl pH 7.5, 100 mM NaCl, 1 mM DTT) to a final salt concentration of 150 mM NaCl and loaded onto a 6 mL RESOURCE Q anion exchange column (Cytiva) pre-equilibrated with low-salt buffer. After washing with 3–5 cv low-salt buffer, proteins were eluted by increasing NaCl concentrations to 1 M in a linear gradient of 60 mL. Peak fractions were pooled and concentrated by ultrafiltration (Vivaspin 30,000 MWCO, Sartorius) before snap freezing in liquid nitrogen.

For condensin I^DC^, the eluate from the calmodulin beads was concentrated by ultrafiltration (Vivaspin 30,000 MWCO, Sartorius) before size-exclusion chromatography (SEC) on a Superose 6 Increase 10/300 column (Cytiva) pre-equilibrated in SEC-buffer (50 mM TRIS-HCl pH 7.5, 200 mM NaCl, 5 % (v/v) glycerol, 1 mM DTT). Peak fractions were pooled and concentrated by ultrafiltration (Vivaspin 30,000 MWCO, Sartorius) before snap freezing in liquid nitrogen.

#### MIX-1–DPY-27_elbow_

DNA fragments encoding *C. elegans* MIX-1 residues 327–860 and wild-type or mutant (E_454_K, K_475_E, K_479_E, E_909_A, K_913_A) DPY-27 residues 441–943 were inserted into pRSET A (ThermoFisher). DPY-27 with an amino-terminal His_8_ tag and untagged MIX-1 were co-expressed in *Escherichia coli* Rosetta (DE3) pLysS (Merck) grown at 37 °C in 2×TY medium. For crosslinking experiments, a fusion construct encoding *C. elegans* DPY-27 residues 441–943 with an amino-terminal His_8_ tag, a 3C protease cleavage sequence as linker, and MIX-1 residues 327–860 with a carboxy-terminal HA_6_ tag was cloned into pRSET A (ThermoFisher) and used for expression in *E. coli* Rosetta (DE3) pLysS grown at 37 °C in 2×TY medium. Cultures were shifted to 18 °C and protein expression was induced by addition of isopropyl-β-D-thiogalactopyranoside (IPTG) to a final concentration of 0.25 mM. Cells were collected 16–18 h post induction by centrifugation, snap frozen in liquid nitrogen, and stored at –80 °C.

Stored cells were thawed and lysed by sonication at 4 °C in lysis buffer (50 mM TRIS-HCl pH 7.5, 300 mM NaCl, 20 mM imidazole, 5 mM β-mercaptoethanol) containing cOm–EDTA. The lysate was cleared by centrifugation at 48,000 ×g for 30 min at 4 °C. The supernatant was incubated with 2 mL Ni Sepharose HP resin (Cytiva) for 2–3 h at 4 °C. The protein-bound resin was packed into a gravity-flow column and washed with 10–20 cv lysis buffer. Bound proteins were eluted with 5–7 cv elution buffer (50 mM TRIS-HCl pH 7.5,100 mM NaCl, 250 mM imidazole). The eluate was dialyzed overnight against IEX buffer (50 mM TRIS-HCl pH 7.5, 100 mM NaCl, 1 mM DTT) at 4° C and loaded onto two 1-mL HiTrap Heparin HP (Cytiva) affinity columns connected in series and pre-equilibrated with IEX buffer. After washing with 3–5 cv IEX buffer, proteins were eluted by increasing NaCl concentrations to 1 M in a linear gradient of 20 cv. Peak fractions were pooled and loaded onto a Superdex 200 increase GL 10/300 column (Cytiva) pre-equilibrated in SEC-buffer (50 mM HEPES-NaOH pH 7.5, 300 mM NaCl, 1 mM DTT). Peak fractions were pooled and concentrated by ultrafiltration (Vivaspin 30,000 MWCO, Sartorius) at 4 °C. The concentrated protein was aliquoted, snap frozen in liquid nitrogen, and stored at –80 °C.

### DPY-21

The coding sequence for *C. elegans* DPY-21 was cloned into pACEBac1, adding a carboxy-terminal TEV-His_8_ tag. A recombinant bacmid was prepared by Tn7 transposition in DH10 Multibac YFP cells^48^ and used for transfection of Sf21 cells. At 72 h post infection, released viruses were harvested and then amplified in Sf21 cells. For protein expression, Sf9 cells were infected with the amplified virus. Cells were harvested 72 h post infection by centrifugation, snap frozen in liquid nitrogen, and stored at –80 °C.

Cells were thawed and lysed by sonication at 4 °C in lysis buffer (50 mM TRIS-HCl pH 7.5, 500 mM NaCl, 20 mM imidazole, 5 mM β-mercaptoethanol) containing cOm–EDTA. The lysate was cleared by centrifugation at 48,000 ×g for 30 min at 4 °C. The supernatant was incubated with 2 mL Ni Sepharose HP resin (Cytiva) for 2–3 h at 4 °C. The protein-bound resin was washed with 10-20 cv of lysis buffer and the proteins were eluted with 5-7 cv elution buffer (50 mM Tris-HCl pH 7.5, 300 mM NaCl, 250 mM imidazole).

Imidazole was removed by overnight dialysis against a SEC buffer (50 mM Tris-HCl pH 7.5, 300 mM NaCl, 1 mM DTT) at 4 °C. The dialyzed eluate was loaded onto a Superose 6 Increase 10/300 column (Cytiva) pre-equilibrated in SEC buffer. Peak fractions were pooled and concentrated by ultrafiltration (Vivaspin 100,000 MWCO, Sartorius). The concentrated protein was aliquoted, snap frozen in liquid nitrogen, and stored at –80 °C.

### DPY-30

*C. elegans* DPY-30 with an amino-terminal His_8_-mCherry-3C tag was cloned into pRSET A (ThermoFisher). The construct was transformed into *E. coli* Rosetta (DE3) pLysS (Merck) grown at 37 °C in 2×TY medium. Protein expression was induced by addition of IPTG to a final concentration of 0.25 mM and cells were shifted to 18 °C. Cells were collected 16–18 h post induction by centrifugation, snap frozen in liquid nitrogen, and stored at –80 °C.

Cells were thawed and lysed by sonication at 4 ^°^C in lysis buffer (50 mM TRIS-HCl pH 7.5, 300 mM NaCl, 20 mM imidazole, 5 mM β-mercaptoethanol) containing cOm–EDTA. The lysate was cleared by centrifugation at 48,000 ×g for 30 min at 4 °C. The supernatant was incubated for 2–3 h with 2 mLNi Sepharose HP (Cytiva) at 4 °C and the protein-bound resin was washed with 10–20 cv lysis buffer. The proteins were eluted with 5–7 cv of elution buffer (50 mM TRIS-HCl pH 7.5, 300 mM NaCl, 250 mM imidazole). The eluate was re-buffered into low-salt buffer (50 mM TRIS-HCl pH 7.5, 100 mM NaCl, 1 mM DTT) by ultrafiltration (Vivaspin 10,000 MWCO, Sartorius) and loaded onto a 6 mL Resource Q ion-exchange column (Cytiva) pre-equilibrated with low-salt buffer. The flowthrough fraction was collected and concentrated by ultrafiltration (Vivaspin 10,000 MWCO, Sartorius). The concentrated protein was aliquoted, snap frozen in liquid nitrogen, and stored at –80 °C.

### SDC-3_C_

The coding sequence for *C. elegans* SDC-3 residues 1,763–2,150 was cloned into a pRSET A (ThermoFisher) vector. Mutants were generated by PCR-based site-directed mutagenesis. The SDC-3_C_ construct with an amino-terminal His_8_-mNeonGreen-3C tag was expressed in *E. coli* Rosetta (DE3) pLysS (Merck) grown at 37 °C in 2×TY medium supplemented with ZnCl_2_ at a final concentration of 25 µM. Protein expression was induced with IPTG at a final concentration of 0.25 mM and cultures were shifted to at 18 °C. Cells were collected 16–18 h post induction by centrifugation, snap frozen in liquid nitrogen, and stored at –80 °C.

Cells were thawed and lysed by sonication at 4 °C in lysis buffer (50 mM TRIS-HCl pH 7.5, 300 mM NaCl, 20 mM imidazole, 0.1 mM ZnCl_2_, 5 mM β-mercaptoethanol) containing cOm–EDTA. The lysate was cleared by centrifugation at 48,000 ×g for 30 min at 4 °C. The supernatant was incubated for 2–3 h with 2 mL Ni Sepharose HP (Cytiva) at 4 °C. The protein-bound resin was washed with 10-20 cv lysis buffer, and proteins were eluted with 5-7 cv elution buffer (50 mM TRIS-HCl pH 7.5,100 mM NaCl, 250 mM imidazole). The eluate was dialyzed overnight against low-salt(Zn^2+^)-buffer (50 mM TRIS-HCl pH 7.5, 100 mM NaCl, 0.1 mM ZnCl_2_, 1 mM DTT) at 4 °C and loaded onto a 6-mL Resource Q ion exchange column (Cytiva) pre-equilibrated with low-salt(Zn^2+^) buffer (50 mM TRIS-HCl pH 7.5, 100 mM NaCl, 0.1 mM ZnCl_2_, 1 mM DTT). After washing with 5 cv low-salt(Zn^2+^) buffer, proteins were eluted by increasing NaCl concentrations to 1 M in a linear gradient of 20 cv. Peak fractions were pooled and concentrated by ultrafiltration (Vivaspin 30,000 MWCO, Sartorius). The concentrated protein was aliquoted, snap frozen in liquid nitrogen, and stored at –80 °C.

For crosslinking experiments, the dialyzed eluate was loaded onto a Superdex 75 pg 16/600 column (Cytiva) pre-equilibrated in HEPES buffer (50 mM HEPES-NaOH pH 7.5, 300 mM NaCl, 0.1 mM ZnCl_2_, 1 mM DTT). Peak fractions were pooled and concentrated by ultrafiltration (Vivaspin 30,000 MWCO, Sartorius). The concentrated protein was aliquoted, snap frozen with liquid nitrogen, and stored at –80 °C.

### Single-molecule DNA loop extrusion assays

Glass slides and λ-phage DNA were prepared and assembled as described^13^. After assembly, flow chambers were flushed with 25 μg/mL streptavidin (Merck) in rinsing buffer (50 mM TRIS-HCl pH 7.5, 20 mM KOAc) and rinsed thoroughly before introducing 1–10 pM biotinylated λ-phage DNA in imaging buffer (50 mM TRIS-HCl pH 7.5, 50 mM KOAc, 2.5 mM MgCl_2_, 5 % (w/v) D-glucose, 1 mM DTT, 500 nM Sytox orange (SxO, ThermoFisher), 40 μg/mL glucose oxidase (Sigma), 15 μg/mL catalase (Sigma), 2 mM Trolox (Sigma), 0.5 mg/mL BSA (NEB)) with a continuous flow of 6–8 μL/min using a PHD2000 syringe pump (Harvard Apparatus).

Unbound DNA molecules were flushed out with imaging buffer before introducing one chamber volume (∼20 μL) of 2–8 nM purified condensin and 1 mM ATP in imaging buffer. For loop extrusion rate analyses, condensin protein concentrations were titrated down to 1–5 nM to avoid multiplex events on the same DNA molecule. For stretching extruded DNA into loops, the flow rate was increased to 50-80 μL/min. For Condensin I^DC^ + SDC-3_C_ experiments, proteins were premixed at molar ratios of 1:1–2 in imaging buffer on ice.

Images were acquired on a THUNDER Imager wide-field microscope (Leica) equipped with an AM TIRF MC module, an iXon Ultra 897EMCCD camera (Andor), and a 100× objective (Leica) at 100-ms exposures using a 532-nm excitation laser.

Imaging data was processed in FIJI with a custom script as described^13,49^.

### Electrophoretic Mobility Shift Assay (EMSA)

The 6-FAM labeled 51-bp dsDNA substrate was prepared by annealing two complementary DNA oligos (Merck, 5’-6-FAM-TATTTTCTTG TTCTTTTACA TCAACACAAT GTGACCATTA CCTGGTTATC A-3’; 5’-TGATAACCAG GTAATGGTCA CATTGTGTTG ATGTAAAAGA ACAAGAAAAT A-3’) for *np1*; (5’-6-FAM-TCGCGAAGGG AGGTGTACCT AGTCTCGTGG ATAAAATATT TGGACAAGGG G-3’; 5’-CCCCTTGTCC AAATATTTTA TCCACGAGAC TAGGTACACC TCCCTTCGCG A-3’) for *rex-32*; in annealing buffer (50 mM TRIS-HCl pH 7.5, 50 mM NaCl) at a concentration of 50 µM using a temperature gradient of 0.1 °C·s^−1^ from 95 °C to 4 °C. The EMSA reaction was prepared with a constant 51-bp dsDNA concentration of 10 nM and increasing concentrations of purified protein in binding buffer (50 mM TRIS-HCl pH 7.5, 50 mM KCl, 125 mM NaCl, 5 mM MgCl_2_, 5 % (v/v) glycerol, 1 mM DTT). After 10 min incubation on ice, free DNA and DNA-protein complexes were resolved by electrophoresis at 4 V·cm^−1^ at 4 °C for 1.5 h in 0.75 % (w/v) TAE-agarose gels. 6-FAM-labeled dsDNA was detected on a Typhoon FLA 9500 laser scanner (GE Healthcare) using a 473-nm excitation laser and a 510-nm LP filter.

### ATP hydrolysis assays

A 6.4-kbp plasmid was relaxed with *E. coli* topoisomerase I (NEB) and purified by phenol/chloroform extraction followed by ethanol precipitation. ATP hydrolysis reactions (10 μL total volume) were set up with 0.5 μM condensin I or I^DC^ in the presence or absence of 25 nM relaxed circular 6.4-kbp plasmid DNA in ATPase buffer (40 mM TRIS-HCl pH 7.5, 125 mM NaCl, 10 % (v/v) glycerol, 5 mM MgCl_2_,). ATP hydrolysis reactions were equilibrated at room temperature for 10 min, started by addition of ATP to final concentrations of 5 mM ATP and 50 nM [α-^32^P]-ATP (Hartmann Analytic), and incubated at room temperature. A volume of 1 μL was spotted onto PEI cellulose F TLC plates (Merck) every 4 min for a total duration of 20 min. The reaction products were separated in 1 M LiCl, 2 M formic acid and signals were detected via a phosphor storage screen scanned on a Typhoon FLA9500 (Cytiva). The ATP hydrolysis rate was calculated by a fit of the ADP/ATP ratios in the linear range of the reaction.

### Microscopy of DPY-27::mStayGold

Strains *dpy-27_wt_* (PMW1268), *him-8(ubs86)* (PMW1329), *his-72::mCherry* (PMW1365), *dpy-27_EKKEKKEEAA_* (PMW1373), *sdc-3_YRAE_* (PMW1337), and *sdc-3*_Δ *2078-2150*_ (PMW1431) (Supplementary Table 1), all expressing endogenously mStayGold-tagged DPY-27, were imaged at embryonic and L1 larval stages using a Nikon Ti2 Yokogawa W1 spinning disk confocal microscope. Image stacks were denoised using Noise2Void (https://github.com/CellFateNucOrg/noise2void) and subsequently maximum intensity projected.

### Immunoprecipitation

Immunoprecipitations of HA_6_-tagged condensin I^DC^ with SDC proteins were performed by mixing proteins at 1 µM concentration in reaction buffer (50 mM TRIS-HCl pH 7.5, 150 mM KOAc, 0.01 % (v/v) Tween-20, 1 mM DTT) followed by 15 min incubation on ice. HA_6_-tagged Condensin I^DC^ was immunoprecipitated by addition of 12.5 μL protein A Dynabeads (ThermoFisher) pre-charged with 3 μg 12CA5 anti-HA monoclonal antibody followed by 40 min incubation on ice. Dynabeads were washed three times with 100 μL reaction buffer. Proteins were eluted by addition of SDS loading buffer (50 mM TRIS-HCl pH 7.5, 1 % (w/v) SDS, 0.1 % (w/v) bromophenol blue, 10 % (v/v) glycerol and 2 % (v/v) β-mercaptoethanol) and heating to 95 °C for 5 min. Eluates were resolved by SDS PAGE.

Immunoprecipitations of mNeon-tagged SDC-3_C_, wild-type and mutant (Y_1890_A, R_1898_E), with MIX-1–DPY-27_elbow_ dimers, wild-type or interface mutant were performed by mixing proteins at 1 µM concentration in reaction buffer (50 mM TRIS-HCl pH 7.5, 150 mM NaCl, 1 mM DTT, 0,1 mM ZnCl_2_, 0.0 2% (v/v) Tween) followed by 15 min incubation on ice. mNeon-tagged SDC-3_C_ was immunoprecipitated by addition of 20 µL mNeonGreen-Trap magnetic agarose (Chromotek) followed by 1 h incubation at 4 °C. Magnetic agarose beads were washed three times with 100 μL reaction buffer. Proteins were eluted by addition of SDS loading buffer and heating to 95 °C for 5 min and then resolved by SDS PAGE.

### Analytical size exclusion chromatography

Condensin I^DC^ and SDC-3_C_ constructs were mixed at an equimolar ratio (2 µM) in a total volume of 70 µL 50 mM TRIS-HCl pH 7.5, 200 mM NaCl, 0.01 mM ZnCl_2_, 1 mM DTT. After incubation on ice for 15 min, samples were loaded onto a Superose 6 Increase 3.2/300 column (Cytiva) pre-equilibrated in the same buffer at a flow rate of 0.1 mL·min^−1^. Fractions of 200 μL were collected and analyzed by SDS PAGE and Coomassie Blue staining.

MIX-1-DPY-27 ‘elbow’ dimer and SDC-3_C_ constructs were mixed at an equimolar ratio (2 µM) in a total volume of 70 µL 50 mM TRIS-HCl, pH 7.5, 150 mM NaCl, 0.01 mM ZnCl_2_, 1 mM DTT. After incubation on ice for 15 min, samples were loaded onto a Superdex 200 Increase 3.2/300 column (Cytiva) pre-equilibrated in the same buffer at a flow rate of 0.01 mL·min^−1^. Fractions of 200 μL were collected and analyzed by SDS PAGE and Coomassie Blue staining.

### Crosslinking mass spectrometry

For crosslinking, purified proteins were diluted in crosslinking buffer (50 mM HEPES-NaOH pH 7.5, 150 mM NaCl, 1 mM DTT) to a final concentration of 2 µM. Bis(sulfosuccinimidyl)suberate (BS3, Pierce) was added to a final concentration of 0.5 mM and the reaction was quenched by addition of TRIS-HCl pH 7.5 to a final concentration of 100 mM after incubation for one hour on ice. Crosslinked samples were fractionated by SDS-PAGE, gels stained with Coomassie brilliant blue, and visible bands corresponding to a molecular weight greater than those of the non-crosslinked condensin subunits were excised.

Gel slices were destained using sequential treatments with 50 % (v/v) ethanol, 5 mM NH₄HCO₃ and 10 mM NH₄HCO and cysteine residues were reduced and alkylated, followed by tryptic in-gel digestion as described previously^50^. Peptide mixtures were separated on a Vanquish Neo system (ThermoFisher; precolumn: µPAC trapping, analytical column: 50cm µPAC Neo) using binary solvent systems consisting of 0.1 % (v/v) formic acid (solvent A) and 80 % (v/v) acetonitrile, 0.1 % (v/v) formic acid (solvent B) with a flow rate of 0.25 μl·min^−1^. Peptides were separated with a gradient of 2 to 22.5 % B in 44 min, followed by 22.5 to 45 % B in 16 min, and 99 % B for 9 min. Eluted peptides were directly injected into an Orbitrap Ascend mass spectrometer (ThermoFisher) operated in a data-dependent acquisition mode.

MS1 scans were acquired in a scan range of m/z 380 to 1,400 at a resolution of 60,000 (at m/z 400) with an automatic gain control (AGC) of 4×10^5^ ions and a maximum fill time of 123 ms. The most intense at least triply charged precursor ions (cycle time scheduled) were selected for stepped higher-energy collisional dissociation (HCD; 24,28,32%NCE). Additional filters for selection of precursors were: 60 s exclusion time, 5×10^4^ min. intensity, monoisotopic peak selection. MS2 data were acquired using scan range mode “Auto” (i.e. the last mass is automatically set to (precursor_mz + 0.8) × charge × 1.02 + 20.0; the first mass is set such as to satisfy the 5–10–15 rule) at a resolution of 30,000, AGC of 1×10^5^ ions and a maximum fill time of 70 ms.

The resulting raw files were converted to mgf format using msconvert (parameters: --filter “peakPicking true 2-” --filter “msLevel 2-” --filter “titleMaker …”) and recalibrated using the median mass shift from an initial linear peptide search using Xi1.8.7^51,52^. Main search was performed using the same Xi installation with final FDR filtering to 1 %. Result files were parsed with CroCo^53^ and subsets of crosslinked residue pairs identified in all three replicates were generated for visualization after applying an additional score-based filter (score > 12). Circos plots were generated using the pyCirclize module.

### SEC-MALS

Protein samples were diluted to a concentration of 1 mg·mL^−1^ in 25 mM TRIS-HCl pH 7.5, 200 mM NaCl, 5 % (v/v) glycerol. A sample volume of 10 µL was resolved over a Superdex 6 Increase 3.2/300 GL column (Cytiva) column at a flow rate of 0.1 mL·min^−1^, equilibrated in the same buffer. Light scattering and refractive index data were collected on microDAWN 3 and microOptilab detectors (Wyatt). Bovine serum albumin (BSA) was used for system normalization and detector alignment.

### Mass photometry

Samples were diluted immediately prior to applying to the mass photometer a concentration of 400 nM in 50 mM TRIS-HCl pH 7.5, 200 mM NaCl and then further diluted to a final concentration of 10 nM upon loading onto glass coverslips on a TwoMP mass photometer (Refeyn). Videos were recorded for the duration of 1 min using AcquireMP v2.4.0. Data were analyzed with DiscoverMP v2.4.0 (Refeyn). Mass calibration curves were recorded with BSA, immunoglobulin G, and thyroglobulin.

### Cryo-EM sample preparation, data collection and processing

Purified *C. elegans* condensin I^DC^(EQ) and SDC-3_C_ were diluted to a final concentration of 1 µM in 50 mM TRIS-HCl pH 7.5, 75 mM NaCl, 5 mM MgCl_2_, 1 mM DTT, 1 mM ATP. After 10 min incubation, n-Octyl-β-D-glucopyranosid (CN23, Roth) was added to a final concentration of 0.05 % (v/v).

A volume of 4 μL was applied onto a 200 R2/2 mesh grid (Quantifoil) after plasma cleaning in a QT GloQube PLUS glow discharge system (Quorum). Plunge freezing was carried out at 4 °C and 100 % humidity on a FEI Vitrobot Mark IV (ThermoFisher) with a wait time of 2 min, blot force 3, and blot time of 1 s with Whatman blotting paper (Cytiva).

Micrographs were collected on a Titan Krios G3 electron microscope (FEI) equipped with a Falcon 4i direct electron detector (ThermoFischer) in two sessions using EPU 3 software (ThermoFisher). A total of 14,940 micrographs (7,221 and 7,719 per session) were collected in counting mode with a total dose of 70 electrons per Å^2^ during an exposure time of 6.35 s, dose fractionated into 1,953 movie frames at defocus ranges of 1.2–3.0 μm. A magnification of 130,000× resulted in a physical pixel size of 0.946 Å.

Cryo-EM data processing was carried out using CryoSPARC v4.7.0^54^. Supplementary Fig. 3,4 show an overview of the cryo-EM processing scheme used for analyzing the datasets.

### Model building and data presentation

As a starting points for model building of the monomeric condensin I^DC^ core complex, AlphaFold models of the individual condensin I^DC^ subunits were used (AF-G5EFJ4-F1, AF-G5EGE9-F1, AF-P48996-F1, AF-Q9U2M1-F1 and AF-Q09591-F1 respectively)^55^. The models were segmented and docked into the monomeric core map using ChimeraX version 1.9^56^ and flexibly fitted in Coot version 0.9.8.94^57^ with positional and secondary structure restraints. The fitted model was manually rebuilt, extended and iteratively improved using Coot, ISOLDE^58^, and real-space refinement in PHENIX version 1.21.2^59^.

For the model of MIX-1–DPY-27_elbow_ bound to SDC-3_C_, an AlphaFold3 prediction^23^ was used as a starting point for model building. The model was manually docked and flexibly fitted into the MIX-1–DPY-27_elbow_–SDC-3_C_ low-resolution map using Coot with positional and secondary structure restraints. The fitted model was manually adjusted and iteratively improved in Coot using the AlphaFold3 prediction as a reference. The model was finalized by map-independent geometry minimization in PHENIX. The CryoEM data collection, processing, and refinement statistics are summarized in Table 1. CryoEM data and structure figures were prepared with the PyMOL Molecular Graphics System version 3.1, Schrödinger, LLC, UCSF ChimeraX^56^, and APBS^60^.

## Data availability

EM density maps have been deposited in the EMDB under accession numbers 56463, 56464, and 56465. Atom coordinates have been deposited in the PDB under accession numbers 9TZA and 9TZB. Mass spectrometry proteomics data have been deposited to the ProteomeXchange Consortium via the PRIDE partner repository^61^ with dataset identifiers PXD073660 and 10.6019/PXD073660. The deposited data will be available as of the date of publication. All other data will be available upon request. Any additional information required to reanalyze the data reported in this paper will be available from the lead contact upon request.

## Acknowledgements

We are grateful to Valeriia Volodkina, Abdulaziz Jaber, and Sol Bravo for technical support. We thank Christian Kraft and Bettina Böttcher for advice and support with cryo-EM data acquisition, Karine Lapouge and Kim Remans of the EMBL Heidelberg Protein Expression and Purification Core Facility for help with mass photometry, and the members of the Meister and Haering groups for discussions. A.V.G. gratefully acknowledges support from a postdoctoral fellowship of the Alexander von Humboldt Foundation. The work was funded by the Deutsche Forschungsgemeinschaft (DFG, German Research Foundation, grant HA5853/4-1 to C.H.H.) and the Swiss National Foundation (grants SNF 310030_212472, 310030E_183425, 310030E_186573 to P.M.). Microscopy was carried out in the Cryo-EM and Biocenter Imaging Core Facilities of the Julius-Maximilians-Universität Würzburg, funded by the Deutsche Forschungsgemeinschaft (DFG, German Research Foundation) – INST 93/903-1 #359471283, INST 93/1042-1 #456578072, INST 93/1143-1 #525040890, and INST 93/1114-1 #512024179. SEC-MALS equipment was funded by the Deutsche Forschungsgemeinschaft (DFG, German Research Foundation) – INST 93/1146-1 #529276114. Mass spectrometry equipment was funded by the Deutsche Forschungsgemeinschaft (DFG, German Research Foundation) – project #540643320.

## Author contributions

I.S. and B.W. generated protein expression constructs; A.V.G. and G.A. expressed and purified proteins, performed DNA and protein binding assays; A.V.G. performed and analyzed DNA loop extrusion assays; D.B. and S.E. generated and imaged *C. elegans* strains and performed image data analyses; A.V.G. performed cryo-EM experiments and data analysis; M.H. built structure models; G.A., J.B., and B.W. performed crosslinking mass spectrometry experiments and data analysis; P.M. and C.H.H. acquired funding and supervised the project; A.V.G., G.A., P.M., and C.H.H. wrote the manuscript with feedback from all authors.

## Competing interests

The authors declare no competing interests.

## Supplementary Data

**Supplementary Figure 1 |** DNA loops extrusion and ATP hydrolysis activities of *C. elegans* condensin I^DC^ and I complexes

**Supplementary Figure 2 |** SDC-3_C_ binds the MIX-1–DPY-27 elbow of Condensin I^DC^ without altering the biochemical properties of the complex

**Supplementary Figure 3 |** Cryo-EM model generation

**Supplementary Figure 4 |** Cryo-EM model evaluation

**Supplementary Figure 5 |** Phenotypic characterization of mutations that disrupt the DPY-27–SDC-3_C_ interface

**Supplementary Figure 6 |** Structural characterization of the condensin I^DC^–SDC-3_C_ complex

**Supplementary Figure 7 |** Unique structural features of the condensin I^DC^ kleisin subunit

## Supplementary Data

**Supplementary Table 1 |** Guide and repair templates for *C. elegans* strain generation

**Supplementary Table 2 |** *C. elegans* strains

**Supplementary Table 3 |** Crosslinking mass spectrometry data condensin holocomplex–SDC-3_C_

**Supplementary Table 4 |** Crosslinking mass spectrometry data MIX-1–DPY-27_elbow_–SDC-3_C_

## Notes

### Competing Interest Statement

The authors have declared no competing interest.

