## Supplementary Figures for "Structural Basis of Condensin Recruitment for X Chromosome Repression"

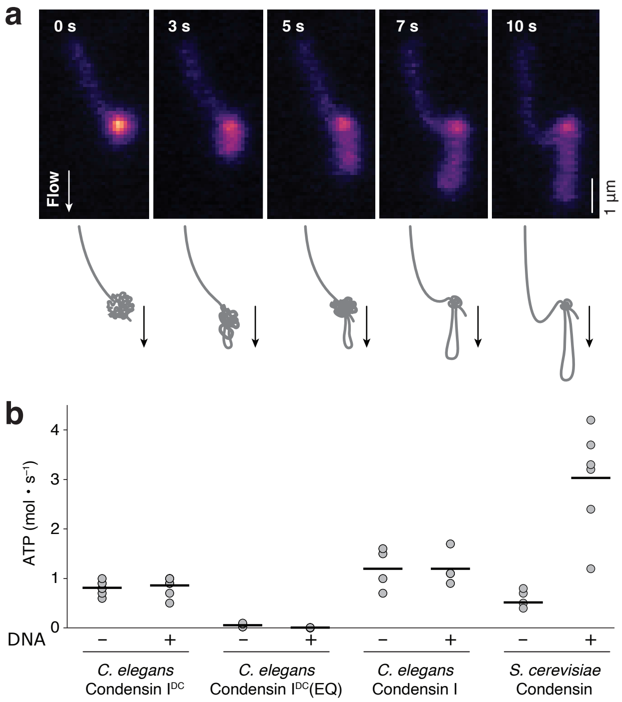


**Supplementary Figure 1 | DNA loops extrusion and ATP hydrolysis activities of *C. elegans* condensin I$^{DC}$ and I complexes**

**a**, Still images from a single-molecule DNA loop extrusion movie using increased flow rates to stretch the extruded DNA into an extended loop structure. **b**, ATP hydrolysis rates of wild-type or a Walker B MIX-1–DPY-27 double mutant (EQ) version of purified *C. elegans* condensin I$^{DC}$ compared to *C. elegans* condensin I or *S. cerevisiae* condensin (0.5 µM) in the absence or presence of DNA (25 nM relaxed circular 6.4-kbp plasmid). Horizontal lines indicate mean values.

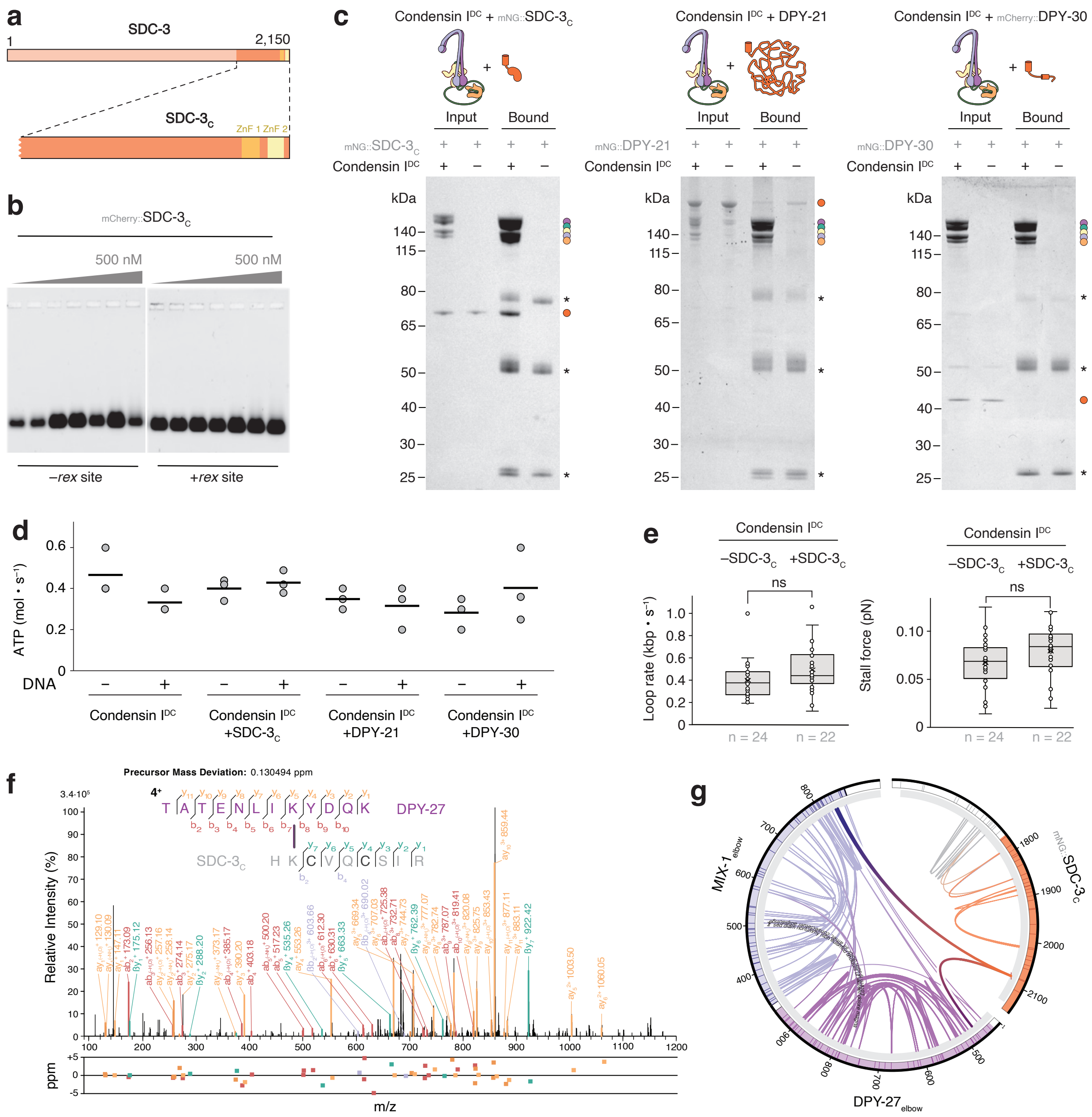

**Supplementary Figure 2 | SDC-$3_C$ binds the MIX-1–DPY-27 elbow of Condensin $I^{DC}$ without altering the biochemical properties of the complex**

**a**, A schematic model of SDC-3 highlights the location of its two zinc finger motifs within its carboxy-terminal domain. **b**, Electrophoretic mobility shift assay (EMSA) of 51-bp dsDNA substrates (10 nM) with or without a *rex* binding site and increasing amounts of mCherry::SDC-$3_C$. **c**, Coomassie Blue-stained SDS PAGE of input and bound fractions of immunoprecipitation reactions against an HA tag fused to the DPY-26 subunit of condensin $I^{DC}$ complexes and SDC-$3_C$ (left), full-length DPY-21 (center), or full-length DPY-30 (right). **d**, ATP hydrolysis rates of condensin $I^{DC}$ (0.5 μM) alone or in the presence of equimolar concentrations of SDC-$3_C$, DPY-21, or DPY-30 in the absence or presence of DNA (25 nM relaxed circular 6.4-kbp plasmid). Horizontal lines indicate mean values. **e**, DNA loop extrusion rates (left) and stall forces (right) of condensin $I^{DC}$ in absence or presence of SDC-$3_C$ (ns = non-significant; $p > 0.05$ in Welch's two-sample t-test). **f**, Mass spec profile of the DPY-27–SDC$3_C$ crosslink identified with the condensin holocomplex. **g**, Circos plot of intra- (thin lines) or inter-molecular (thick lines) crosslinks identified in the MIX-1–DPY-$27_{elbow}$ dimer bound to SDC-$3_C$.

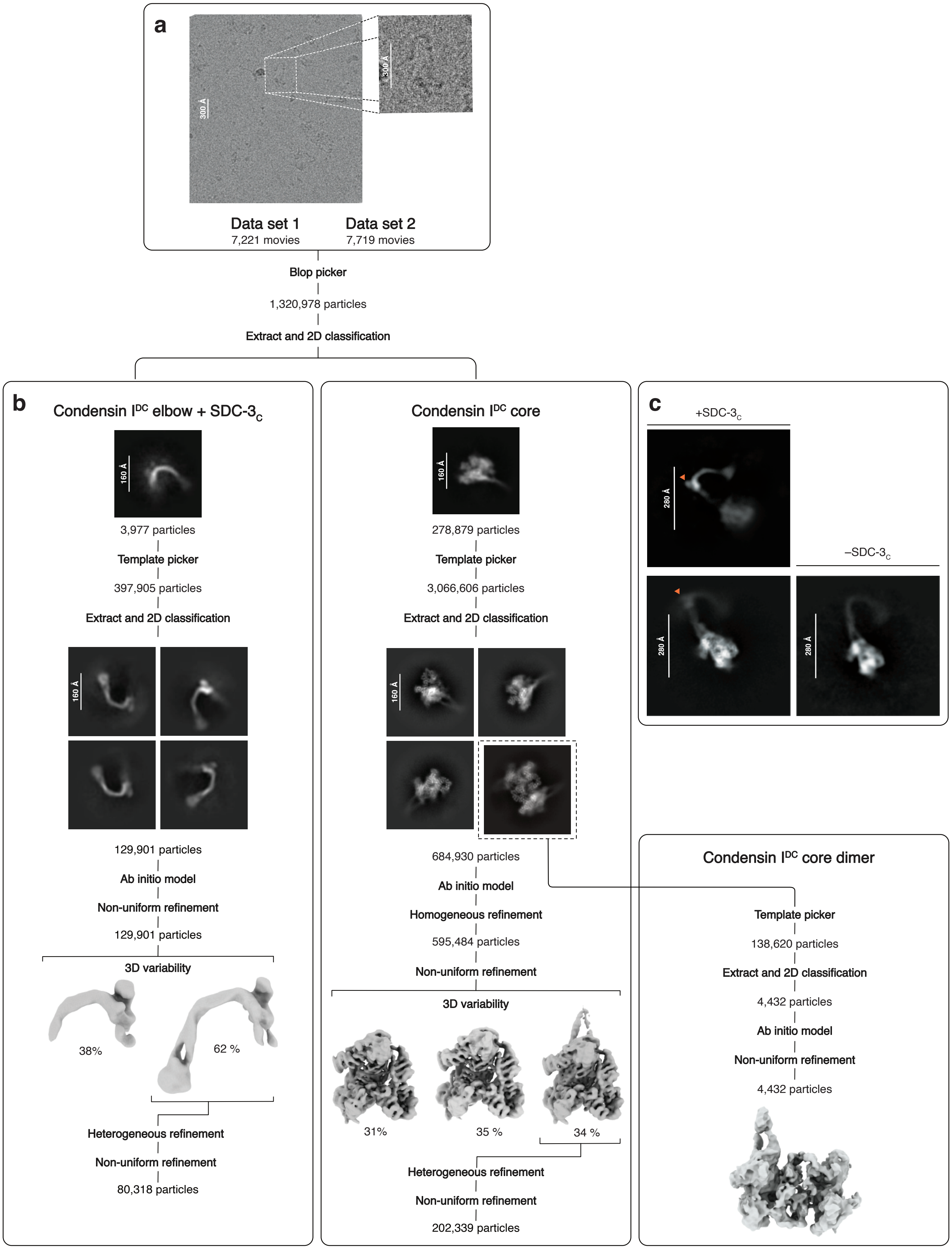


**Supplementary Figure 3 I Cryo-EM model generation**

**a**, Representative micrograph of *C. elegans* condensin I$^{DC}$ complexes bound by SDC-3$_C$. **b**, Workflow of initial data processing of MIX-1/DPY-27elbow bound to SDC-3$_C$ (left) or the condensin IDC core monomer (center) or dimer (right) subcomplexes, showing representative 2D classes and 3D-refined maps. **c**, Representative 2D classes of condensin I$^{DC}$ holocomplexes in the presence (left) or absence (right) of SDC-3$_C$. Arrows indicate extra density of SDC-3$_C$ bound to the elbow region of the MIX-1–DPY-27 coiled coil.

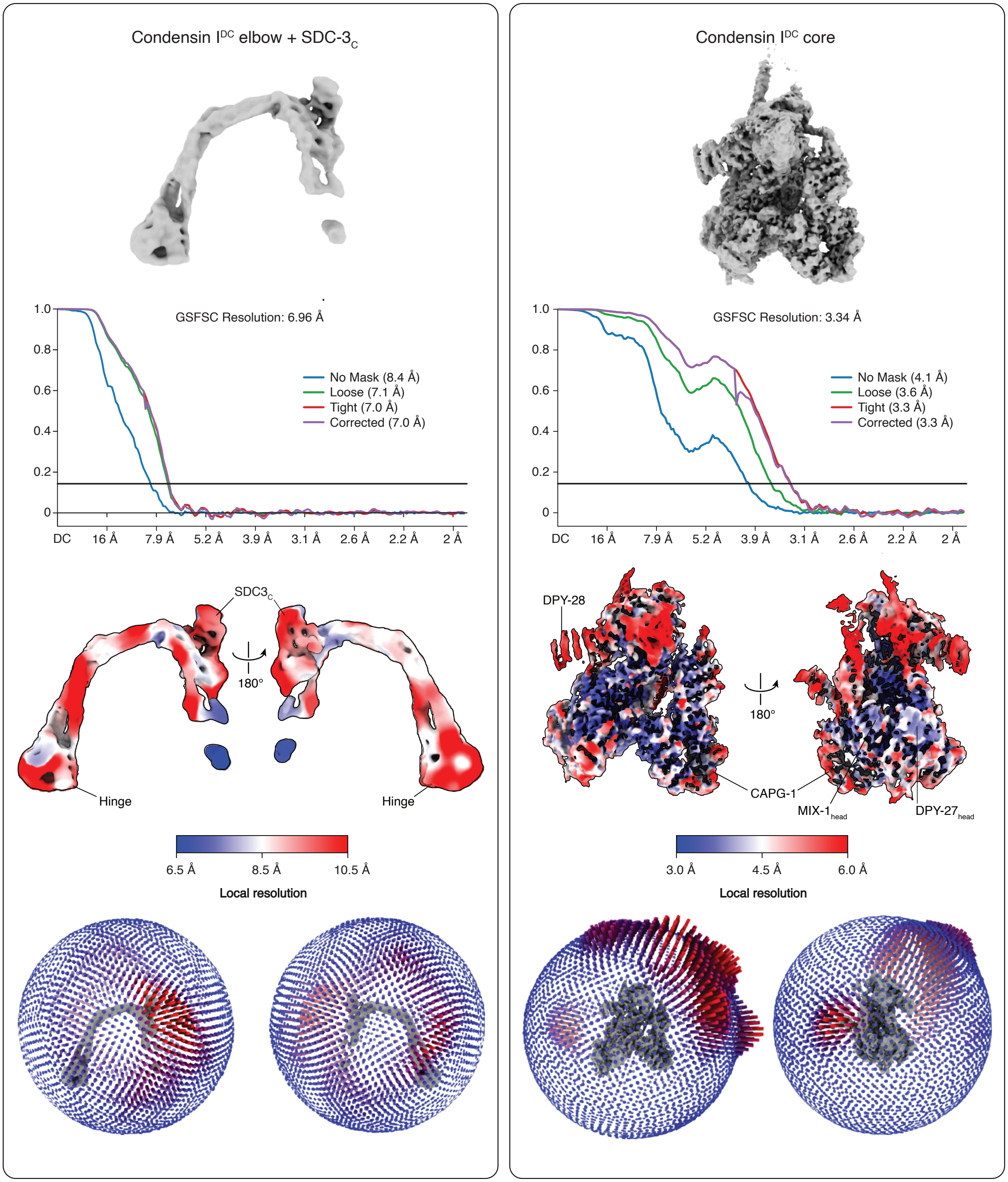


**Supplementary Figure 4 | Cryo-EM model evaluation.**

FSC curves, local resolution maps, and angular distribution plots of the processed maps of MIX-1–DPY-27$_{elbow}$ bound to SDC-3$_C$ (left) or the condensin I$^{DC}$ core monomer (right) resolved to nominal resolutions of 6.96 Å or 3.34 Å, respectively.

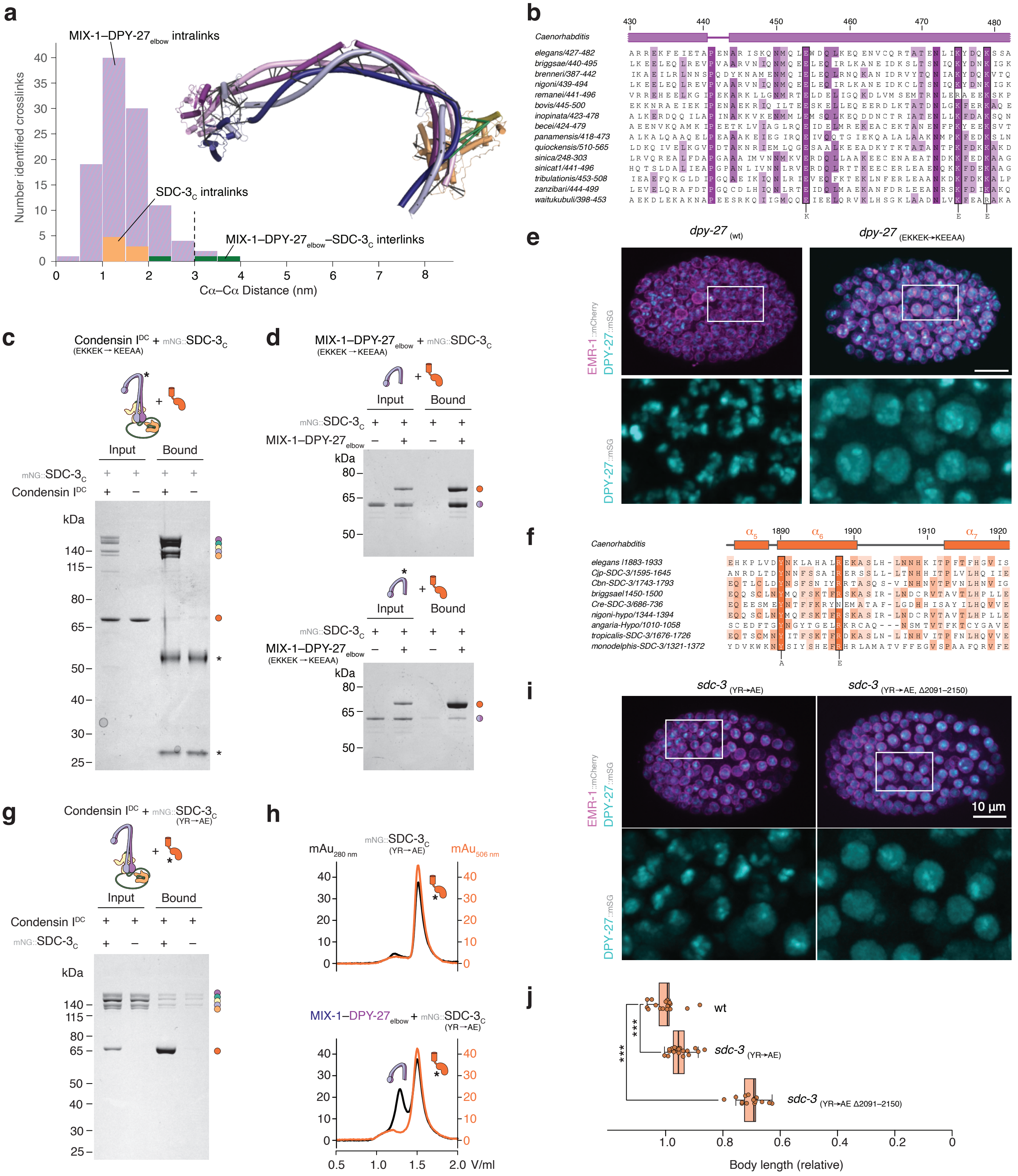

a
MIX-1–DPY-27elbow intralinks
SDC-3C intralinks
MIX-1–DPY-27elbow–SDC-3C interlinks
Number identified crosslinks
Cα–Cα Distance (nm)
b
Caenorhabditis
elegans/427-482
briggsae/440-495
brenneri/387-442
nigoni/439-494
remanei/441-496
bovis/445-500
inopinata/423-478
becei/424-479
panamensis/418-473
quiockensis/510-565
sinica/248-303
sinicat1/441-496
tribulationis/453-508
zanzibari/444-499
waitukubuli/398-453
c
Condensin IDC + mNG::SDC-3C (EKKEK → KEEAA)
Input
Bound
mNG::SDC-3C
Condensin IDC
kDa
d
MIX-1–DPY-27elbow + mNG::SDC-3C (EKKEK → KEEAA)
Input
Bound
mNG::SDC-3C
MIX-1–DPY-27elbow
kDa
MIX-1–DPY-27elbow (EKKEK → KEEAA)
e
dpy-27 (wt)
dpy-27 (EKKEK→KEEAA)
EMR-1::mCherry
DPY-27::mSG
f
Caenorhabditis
elegans I1883-1933
Cjp-SDC-3/1595-1645
Cbn-SDC-3/1743-1793
briggsaeI1450-1500
Cre-SDC-3/686-736
nigoni-hypo/1344-1394
angaria-Hypo/1010-1058
tropicalis-SDC-3/1676-1726
monodelphis-SDC-3/1321-1372
g
Condensin IDC + mNG::SDC-3C (YR→AE)
Input
Bound
Condensin IDC
mNG::SDC-3C
kDa
h
mNG::SDC-3C (YR→AE)
mAu280 nm
mAu506 nm
MIX-1–DPY-27elbow + mNG::SDC-3C (YR→AE)
V/ml
i
sdc-3 (YR→AE)
sdc-3 (YR→AE, Δ2091–2150)
EMR-1::mCherry
DPY-27::mSG
10 µm
j
wt
sdc-3 (YR→AE)
sdc-3 (YR→AE Δ2091–2150)
***
Body length (relative)

**Supplementary Figure 5 | Phenotypic characterization of mutations that disrupt the DPY-27–SDC-3$_C$ interface**
**a**, Length distribution of MIX-1–DPY-27$_{elbow}$ intra-crosslinks (blue-purple striped), SDC-3$_C$ intra-crosslinks (orange), or inter-crosslinks between MIX-1–DPY-27$_{elbow}$ and SDC-3$_C$ (green). Intra-crosslinks (dark grey) and inter-crosslinks (green) are mapped onto the cartoon model of the elbow interface. **b**, DPY-27$_{elbow}$ homologous protein sequence alignment from different *Caenorhabditis* species. **c**, Coomassie Blue- stained SDS PAGE of input and bound fractions of immunoprecipitation reactions against an HA tag fused to the DPY-26 subunit of condensin IDC complexes containing the DPY-27 E$_{454}$K, K$_{475}$E, K$_{479}$E, E$_{909}$A, K$_{913}$A mutant and mNeonGreen (mNG)-tagged SDC-3$_C$. Asterisks mark immunoglobulin chains in the bound fractions. **d**, Coomassie Blue-stained SDS PAGE of input and bound fractions of immunoprecipitation reactions against mNG fused to the SDC-3$_C$ and wild-type (top) or interface mutant (bottom) MIX-1–DPY-27$_{elbow}$ dimers and SDC-3$_C$. **e**, Representative confocal microscopy images of late *C. elegans* hermaphrodite embryos expressing mCherry-labeled EMR-1 as marker for the nuclear envelope and wild-type (left) or E$_{454}$K, K$_{475}$E, K$_{479}$E, E$_{909}$A, K$_{913}$A mutant (right) versions of DPY-27 fused to mStayGold. **f**, SDC-3$_C$ homologous protein sequence alignment from different Caenorhabditis species. **g**, Coomassie Blue-stained SDS PAGE of condensin I$^{DC}$ input and bound fractions of immunoprecipiation reactions against the Y$_{1890}$A, R$_{1898}$E interface mutant of mNG::SDC-3$_C$. **h**, Analytical SEC profiles for the Y$_{1890}$A, R$_{1898}$E mutant version of mG::SDC-3$_C$ in the absence (left) or presence (right) of the MIX-1–DPY-27$_{elbow}$ dimer. **i**, Representative confocal microscopy images of late *C. elegans* hermaphrodite embryos expressing mCherry-labeled EMR-1 as marker for the nuclear envelope, DPY-27 fused to mStayGold, and the homozygous *sdc-3* Y$_{1890}$A, R$_{1898}$E mutant version without (left) or with (right) an additional deletion of its zinc finger motifs (Δ2091–2150). **j**, Relative body lengths of homozygous *sdc-3* Y$_{1890}$A, R$_{1898}$E mutants without (strain 1,337) or with (strain 1,370) an additional deletion of its zinc finger motifs (Δ2091–2150) compared to wild-type animals (strain 1,297, from Figure 5D). The box shows the interquartile range (Q1 to Q3) with a line for the median; whiskers show minimum and maximum values within the 1.5× interquartile range (*** $p < 0.001$, t-test or Wilcoxon rank-sum test, depending on normality and variance assumptions, with p-values adjusted for multiple comparisons using the FDR method).

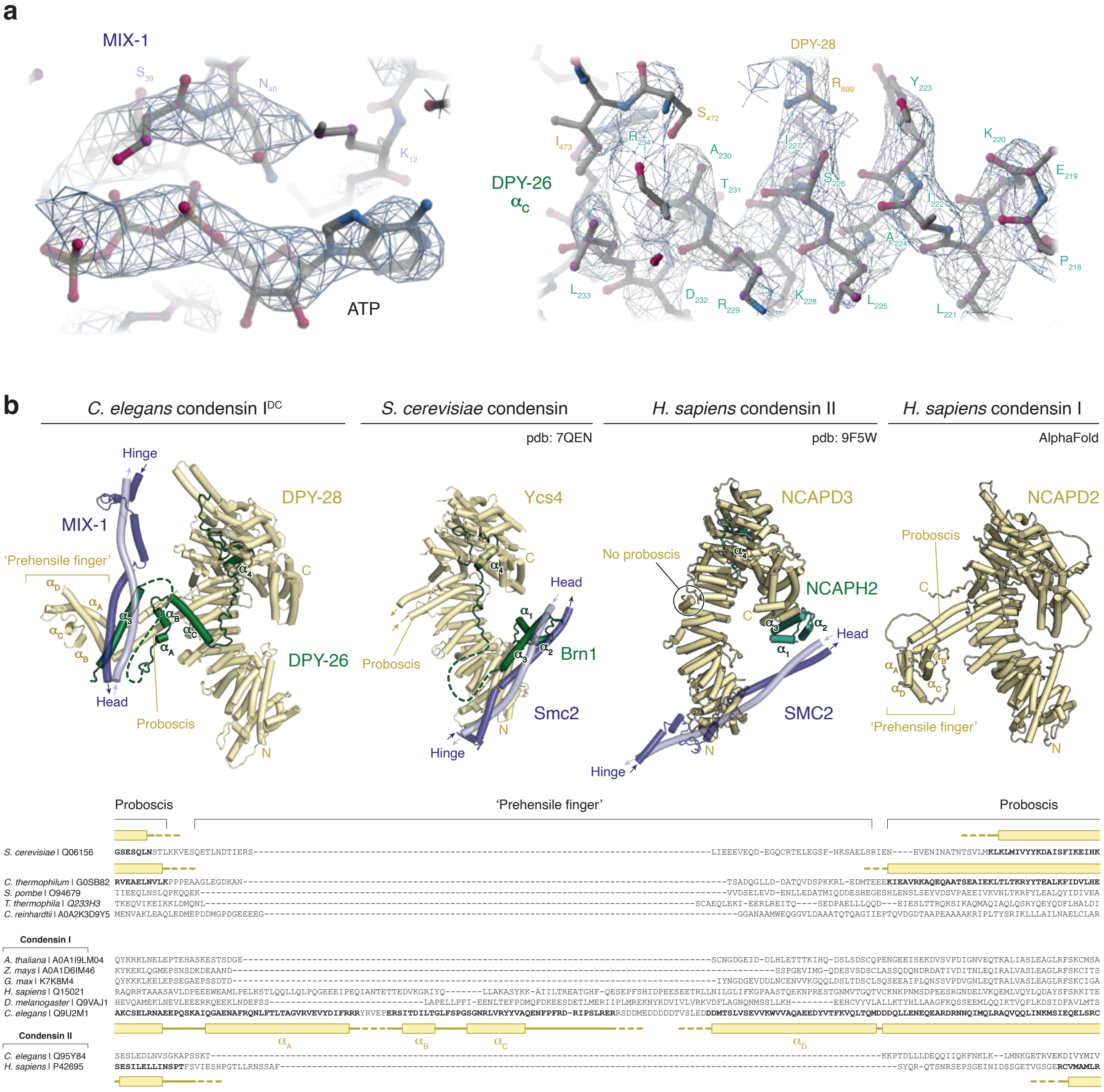


**Supplementary Figure 6 | Structural characterization of the condensin I$^{DC}$–SDC-3$_C$ complex**

**a**, Examples of the electron density maps of ATP bound to the Walker-A motif of MIX-1 (left) and the DPY-26 insertion helix $\alpha_C$ bound to the DPY-28 proboscis (right). **b**, Comparison of HEAT-A (yellow) proboscis structures of *C. elegans* condensin I$^{DC}$, *S. cerevisiae* condensin (pdb: 7QEN), human condensin II (pdb: 9F5W), and human condensin I (predicted by AlphaFold).

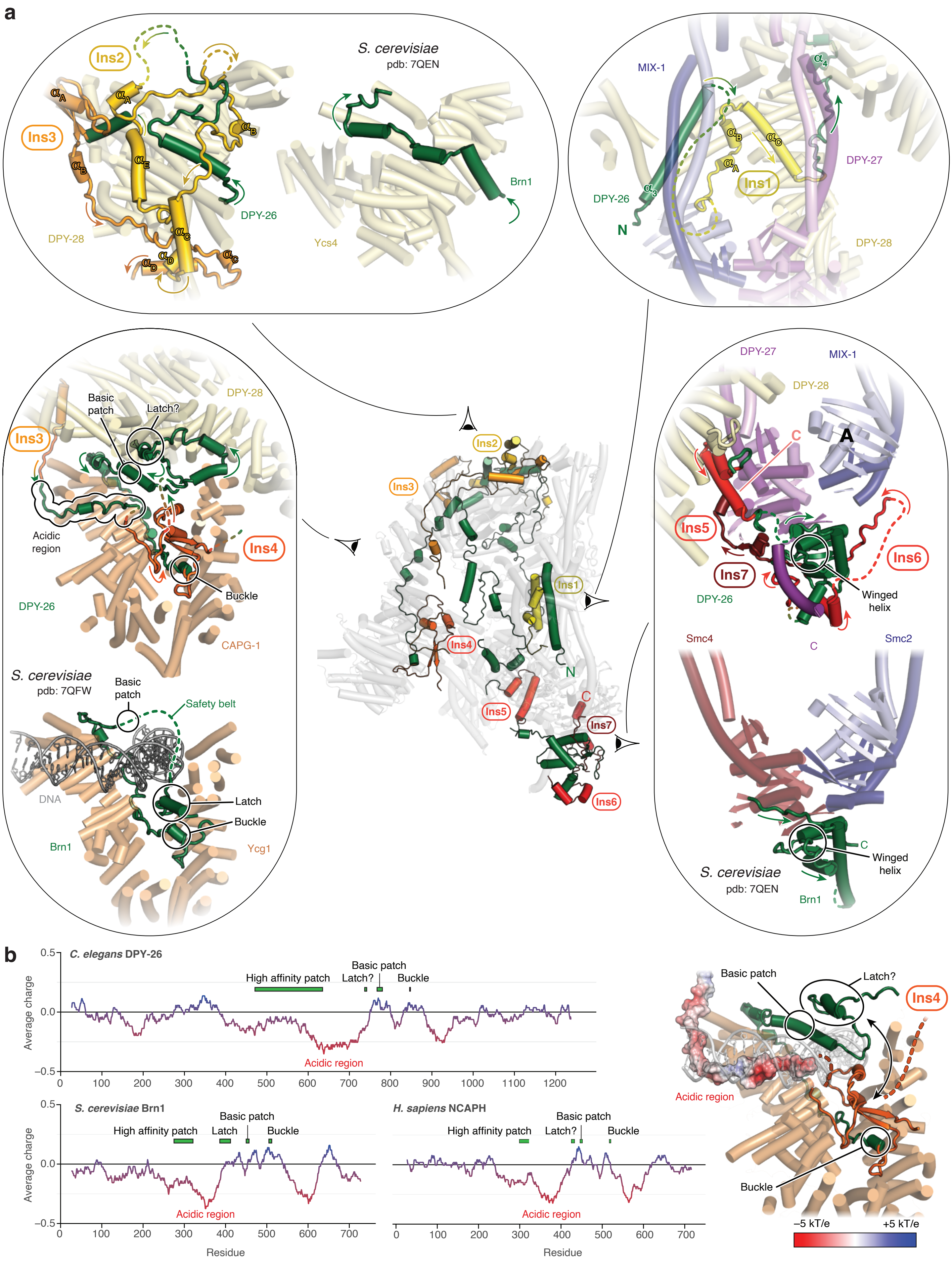

a
S. cerevisiae
pdb: 7QEN
Ins2
Ins3
DPY-26
DPY-28
Brn1
Ycs4
MIX-1
DPY-27
DPY-26
N
Ins1
DPY-28
DPY-28
Basic patch
Latch?
Ins3
Acidic region
Ins4
Buckle
DPY-26
CAPG-1
S. cerevisiae
pdb: 7QFW
Basic patch
Safety belt
DNA
Latch
Buckle
Brn1
Ycg1
Ins2
Ins3
Ins1
Ins4
N
Ins5
C
Ins7
Ins6
DPY-27
MIX-1
DPY-28
C
A
Ins5
Ins6
Ins7
DPY-26
Winged helix
Smc4
C
Smc2
C
Winged helix
S. cerevisiae
pdb: 7QEN
Brn1
b
C. elegans DPY-26
High affinity patch
Basic patch
Latch?
Buckle
Acidic region
Average charge
S. cerevisiae Brn1
High affinity patch
Latch
Basic patch
Buckle
Acidic region
H. sapiens NCAPH
High affinity patch
Latch?
Basic patch
Buckle
Acidic region
Residue
Residue
Basic patch
Latch?
Ins4
Acidic region
Buckle
−5 kT/e
+5 kT/e

**Supplementary Figure 7 | Unique structural features of the condensin I$^{DC}$ kleisin subunit**

**a**, Cartoon model of the path of the kleisin subunit (green) and DPY-26-specific insertions (Ins1–Ins7; from yellow to red). The insets compare the kleisin structures of the HEAT-A high-affinity patch (top left), HEAT-B binding region (left), and κ-SMC binding region (right) to the homologous regions of the *S. cerevisiae* condensin structure (pdb: 7QEN or 7QFW). A separate inset (top right) highlights the HEAT-A proboscis interface. **b**, Calculation of local charge with a sliding window of 51 residues for the *C. elegans* condensin I$^{DC}$, *S. cerevisiae* condensin, and human condensin I kleisin subunits. The cartoon model highlights an acidic region that precedes the safety belt and occupies the HEAT-B site bound by DNA in the homologous *S. cerevisiae* structure (pdb: 7QFW). The electrostatic surface is shown for this part of the kleisin (negative –5 kT/e, red; positive +5 kT/e, blue).
