## Supplementary Table 2 for "Structural Basis of Condensin Recruitment for X Chromosome Repression"

### Supplementary Table 2 | *C. elegans* strains

| Strain name | Genotype |
|---|---|
| PMW1268 | *dpy-27(ubs81[mStayGold]) III; bqSi225[pBN34(unc-119(+) Pemr-1::emr-1::mcherry)] IV* |
| PMW1329 | *him-8(ubs86); bqSi225[pBN34(unc-119(+) Pemr-1::emr-1::mcherry)] IV; dpy-27(ubs81[mStayGold]) III* |
| PMW1337 | *dpy-27(ubs81[mStayGold]) III; bqSi225[pBN34(unc-119(+) Pemr-1::emr-1::mcherry)] IV; sdc-3(ubs52[C-term Halotag] ubs97[Y1890A, R1898E, intron 11 deletion]) V* |
| PMW1365 | *his-72(kog5[his-72::mCherry]); dpy-27(ubs81[mStayGold]) III* |
| PMW1368 | *his-72(kog5[his-72::mCherry]); dpy-27(ubs81[mStayGold]) III; him-8(ubs86); bqSi225[pBN34(unc-119(+))]* |
| PMW1370 | *bqSi142[pBN20(unc-119(+) Pemr-1::emr-1::mCherry)] II; dpy-27(ubs81[mStayGold]) III; bqSi225[pBN34(unc-119(+) Pemr-1::emr-1::mcherry)] IV; sdc-3(ubs52[C-term Halotag] ubs97[Y1890A, R1898E, intron 11 deletion] ubs111[del2091-2150]) V / tmC12 [egl-9(tmIs1197)] V* |
| PMW1373 | *dpy-27(ubs81[mStayGold] ubs105[E454K, K475E, K479E] ubs109[E909A, K913A]) III; bqSi225[pBN34(unc-119(+) Pemr-1::emr-1::mcherry)] IV; sdc-3(ubs52[C-term Halotag]) V* |
| PMW1431 | *dpy-27(ubs81[mStayGold]) III;*<br>*sdc-3(ubs52[C-term Halo]; ubs70[p.2078_2150del) V/tmC12 [egl-9(tmIs1197)] V;*<br>*bqSi142[pBN20(unc-119(+) Pemr-1::emr-1::mCherry)] II* |
